# Rationalizing diverse binding mechanisms to the same protein fold: insights for ligand recognition and biosensor design

**DOI:** 10.1101/2024.03.25.586677

**Authors:** Alison C. Leonard, Anika J. Friedman, Rachel Chayer, Brian M. Petersen, Joel Kaar, Michael R. Shirts, Timothy A. Whitehead

**Affiliations:** Department of Chemical and Biological Engineering, University of Colorado Boulder, Boulder, CO, 80305, USA

**Keywords:** protein-ligand binding, protein biosensors, deep mutational scanning, molecular dynamics, computational protein design, Rosetta

## Abstract

The engineering of novel protein-ligand binding interactions, particularly for complex drug-like molecules, is an unsolved problem which could enable many practical applications of protein biosensors. In this work, we analyzed two engineer ed biosensors, derived from the plant hormone sensor PYR1, to recognize either the agrochemical mandipropamid or the synthetic cannabinoid WIN55,212-2. Using a combination of quantitative deep mutational scanning experiments and molecular dynamics simulations, we demonstrated that mutations at common positions can promote protein-ligand shape complementarity and revealed prominent differences in the electrostatic networks needed to complement diverse ligands. MD simulations indicate that both PYR1 protein-ligand complexes bind a single conformer of their target ligand that is close to the lowest free energy conformer. Computational design using a fixed conformer and rigid body orientation led to new WIN55,212-2 sensors with nanomolar limits of detection. This work reveals mechanisms by which the versatile PYR1 biosensor scaffold can bind diverse ligands. This work also provides computational methods to sample realistic ligand conformers and rigid body alignments that simplify the computational design of biosensors for novel ligands of interest.

## Introduction

The design and engineering of proteins for specific, reversible, and high affinity binding with small molecule ligands remains a grand challenge in biotechnology. Fundamentally, design presents a stringent test for the predictive control of molecular recognition events. Practically, new protein-ligand binders can drive new biosensors where the molecular recognition domain is integrated with or coupled to an output signal ^1^. Functional biosensors can enable a wide range of biotechnologies including recent examples in agrochemical control of plant traits ^2^, real-time analysis of neurotransmitter activity ^3^, and spatiotemporal control of cellular therapies ^4–7^.

New protein-ligand binders have been created by reengineering an existing binding site to recognize different new molecules ^8–13^ or by screening sequence libraries to identify binders ^14^. There have also been several reports of computationally designed protein binders ^15–21^. These computationally designed proteins bind just a handful of ligands that are not fully representative of drug-like molecules which contain many rotatable bonds and multiple functional groups ^1,22^. To inform the engineering and design of new protein biosensors, there is a pressing need to understand protein binding to a broader range of more complex and flexible small molecules.

Many of the above sensors are bespoke designs, where one protein scaffold or fold binds one unique ligand. However, several protein folds have evolved and been engineered to bind diverse ligands with affinity and specificity. For example, the immunoglobulin fold used by antibodies is quite successful in the molecular recognition of both protein and small molecule ligands ^23,24^. Members of both the lipocalin fold and the START superfamily naturally bind, and can further be engineered to bind, a variety of small molecule ligands ^25–30^. Richer information on the sequence, structural, and mechanistic basis of ligand binding may be found by interrogating a sensor family’s recognition of distinct ligands rather than bespoke designs.

The START domain superfamily member PYR1 has recently been engineered to recognize dozens of natural and synthetic cannabinoids, organophosphates, and fungicides with micromolar to picomolar EC50s ^2,13,31^. PYR1, along with its binding partner HAB1, is part of a natural chemically induced dimerization (CID) system utilized by the plant hormone abscisic acid ^32,33^. Engineered PYR1 binds its cognate ligand independently from HAB1, which then enables HAB1 recognition to form a ternary complex (**Figure 1A**) ^34^. This ‘molecular ratchet’ architecture is particularly well-suited for biosensors because the same molecular recognition component can be coupled to many different output signals ^1^.

**Figure 1.**
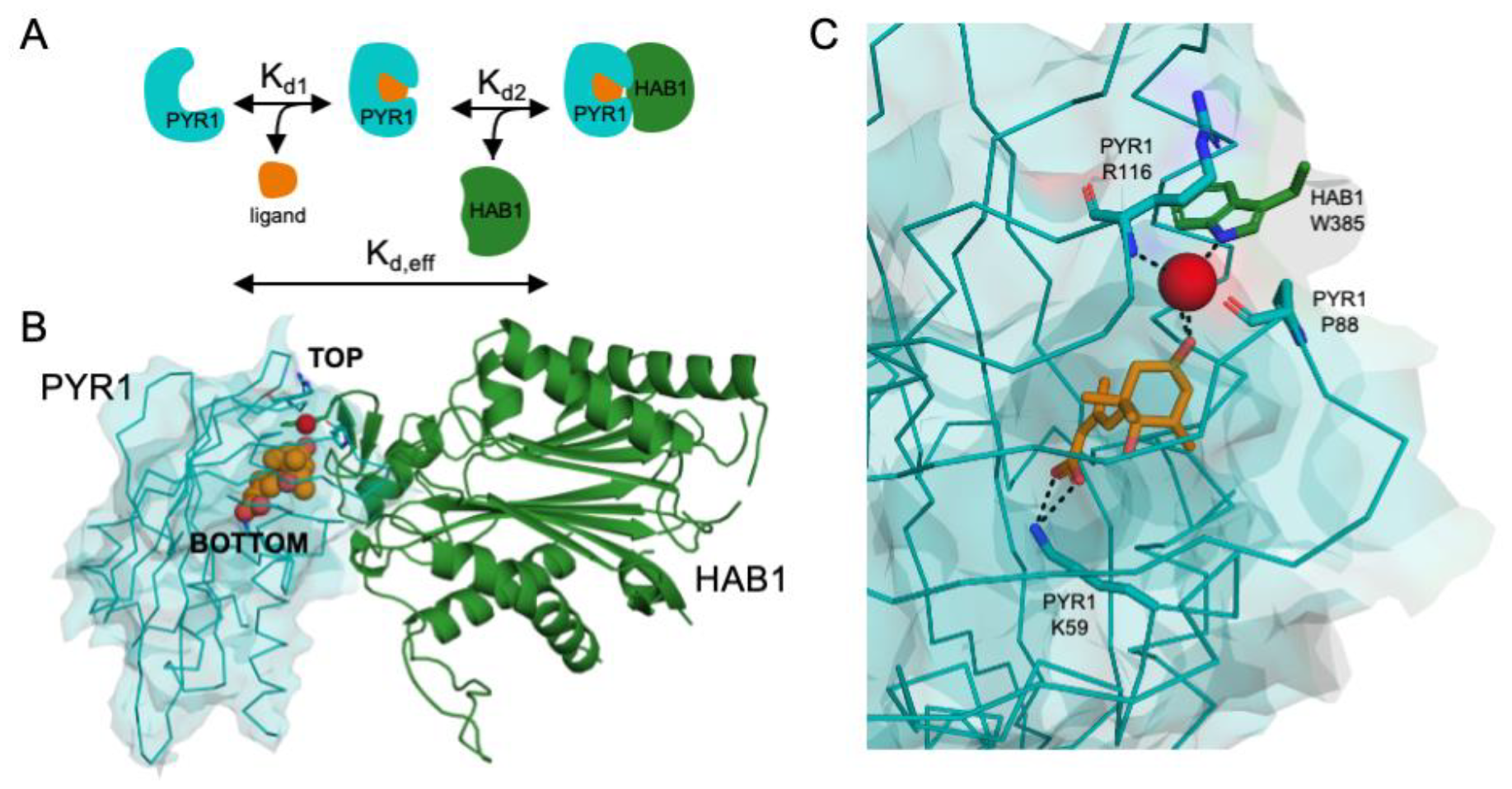
An overview of PYR1-HAB1 binding and structural features. A. Cartoon of two-step binding process in which PYR1 first binds its ligand before dimerizing with HAB1. An effective binding constant K_d,eff_ can describe the overall reaction equilibrium. B. Structure of PYR1 (blue) with abscisic acid (orange) bound, complexed with HAB1 (green) (PDB 3QN1), with the top and bottom of the binding pocket labeled. The bound water coordinating abscisic acid and HAB1 is shown as a red sphere. C. Abscisic acid binding is facilitated by the coordination of a water molecule (red sphere) at the top of the pocket by PYR1 residues P88 and R116, HAB1 W385, and an abscisic acid carbonyl. Abscisic acid is stabilized in the bottom of the pocket by electrostatic complementary between PYR1 K59 and the abscisic acid carboxylate.

Crystal structures of both the wild-type and several engineered sensors all show a similar gate-latch-lock mechanism^13,35^, in which a hydrogen bond acceptor on the bound ligand coordinates a key water molecule at the top of the binding pocket (**Figure 1B**) ^13^. The role of this water molecule appears to be distinct from other waters resolved in the electron density. Despite this common latch mechanism ^36^, it is unclear how this protein binding pocket can be engineered to form the varied molecular interactions that would be needed to bind such diverse chemical ligands with the described high affinity and specificity.

Understanding the sequence, structural, and mechanistic determinants of binding is essential for identifying the key residues that must remain intact for each ligand to bind and allow formation of the ternary complex, as well as which positions are amenable to mutations. Such insight can also help uncover mechanisms for promoting or curtailing ligand recognition, such as the formation of salt bridges or other electrostatic interactions. As water is known to mediate the gate-latch-lock closure mechanism, an analysis of the role of other water molecules present in the electron density of crystal structures will determine other ways in which water may influence the protein sequence, and binding capacity for diverse ligands. Determining the set of ligand conformers that may fit into the binding pocket could uncover metrics for predicting the allowable conformer space, which is critical for protein-ligand binding design.

To understand why engineered PYR1 scaffolds recognize such a diverse array of ligands, we used deep mutational scanning to map the sequence determinants of binding for two engineered biosensors, one sensitive to the agrochemical mandipropamid (PYR1^mandi^) and the other to the synthetic cannabinoid (+)-WIN55,212-2 (PYR1^WIN^). These two sensors differ by amino acid mutations at a total of 7 positions, and the ligands differ by 4 carbons and 6 rotatable bonds ^37^. Mutational landscape maps were complemented with molecular dynamics (MD) simulations to probe the unique importance of the coordinated water molecules of the gate-latch-lock closure mechanism, compared to other present waters, and to visualize the role of salt bridges on mechanisms of binding. For each sensor, we identified critical positions restricted to only 1-2 allowable amino acids and highlighted key differences in the electrostatic interactions required to complement different ligand structural features. This contrasts with positions that exhibited comparably higher residue flexibility in the binding pocket and illustrates, along with MD analysis of conformer preference and experimental characterization of variants with additive mutations, how compatible conformers may have been selected for during directed evolution of the sensor. Using insights from this study, we computationally designed new PYR1 sensors for the synthetic cannabinoid (±)-WIN55,212-2. This detailed study of molecular recognition maps the essential mutations for high-affinity ligand binding in the PYR-HAB biosensor scaffold, highlighting differences in binding principles between dissimilar ligands on a common scaffold and informing the requirements for computational and rational design of biosensors for other diverse ligands.

## Results

### Quantitative binding affinity measurements using deep mutational scanning

We used deep mutational scanning to assess the sequence determinants of ligand recognition for the PYR1^mandi^ and PYR1^WIN^ engineered biosensors (full genotypes listed in bold>Supplemental **Table S1**). For each biosensor we created a single site-saturation mutagenesis library at 27 positions inside and adjacent to the PYR1 pocket. Mutations to either cysteine or stop codons were excluded from analysis. Each library was cloned into a yeast surface display vector and expressed for evaluation by fluorescence activated flow cytometry. To evaluate variant binding, each library was sorted at a constant HAB1^T+^ (a computationally thermostabilized ΔN-HAB1 ^13^) concentration over a range of ligand concentrations, with the top 15-25% of binders collected for each ligand concentration (**Figure 2A**). These sorted populations were then sequenced, with the number of reads in the sorted population compared to the number of reads in a reference population of all displayed protein variants, to calculate the probability that each variant was collected. The effective dissociation constant, K_d,eff_, for each variant can be calculated by maximum likelihood estimation algorithm from MAGMA-seq ^38^. The expression log(Kd,eff,i/Kd,eff,WT) is used to quantify the relative binding of variant i compared to the wild-type sensor.

**Figure 2.**
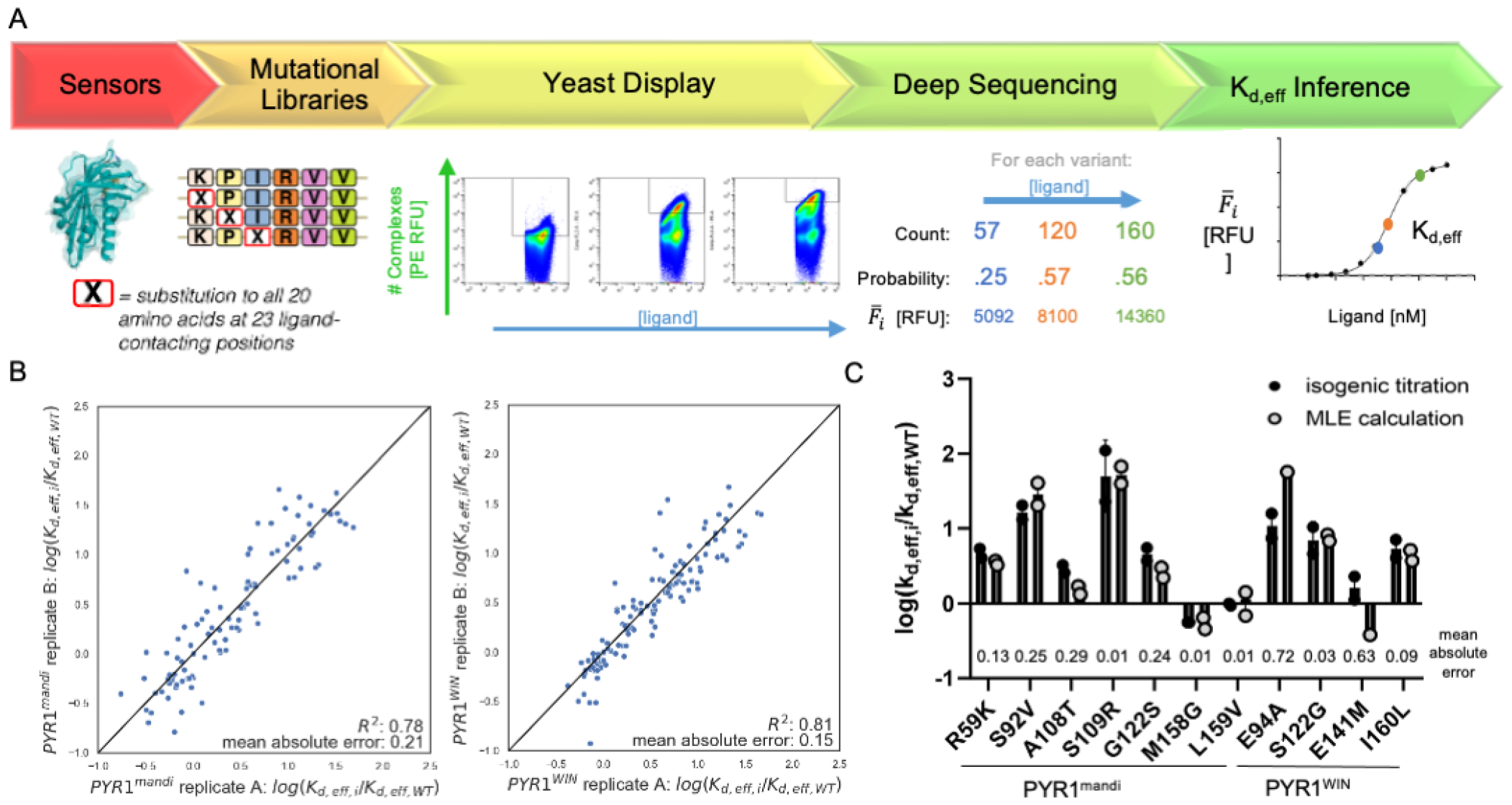
Deep mutational scanning was used to calculate the quantitative effective binding affinities of many biosensor variants in parallel. ***A***. An overview of library sorting and deep mutational scanning to reconstruct binding curves and calculate the effective dissociation constant K_d,eff,i_ for each protein variant. B. Comparison of calculated log(K_d,eff,i_/K_d,eff,WT_) values for replicate sorting experiments for the PYR1^mandi^ and PYR1^WIN^ sensors. C. Comparison of calculated log(K_d,eff,i_/K_d,eff,WT_) values to isogenic titration values for engineered biosensors.

The value of log(Kd,eff,i/Kd,eff,WT) for the PYR1^mandi^ and PYR1^WIN^ mutational variants calculated from deep mutational scanning of library titrations are reported in **supplementary dataset 1**. To assess the internal reproducibility of our protocol, biological replicates of the PYR1^mandi^ and PYR1^WIN^ libraries were sorted on separate days and processed as duplicates. For the PYR1^mandi^ sensor, we observed an R^2^ correlation of 0.78 and mean absolute error (MAE) in log-units of 0.21 (approx. 60% unsigned error in the relative K_d,eff_;

**Figure 2B**). Biological replicates for the PYR1^WIN^ sensor processed on different days have an R^2^ = 0.81 and a mean absolute error of 0.15 in log units (**Figure 2B**). We also performed single variant (isogenic) titrations of 11 PYR1^mandi^ and PYR1^WIN^ mutational variants spanning over a 100-fold range of inferred Kd,eff,i/Kd,eff,WT values. The average MAE was 0.22 log units relative to the inferred values from the population measurements (**Figure 2D-E)**, indicating a similar experimental reproducibility between population and individual measurements. The heatmaps in figures 3 and 4 include a subset of the data encompassing the 23 ligand-facing positions and display the average of two replicates unless a value could be calculated from only one replicate.

**Figure 3.**
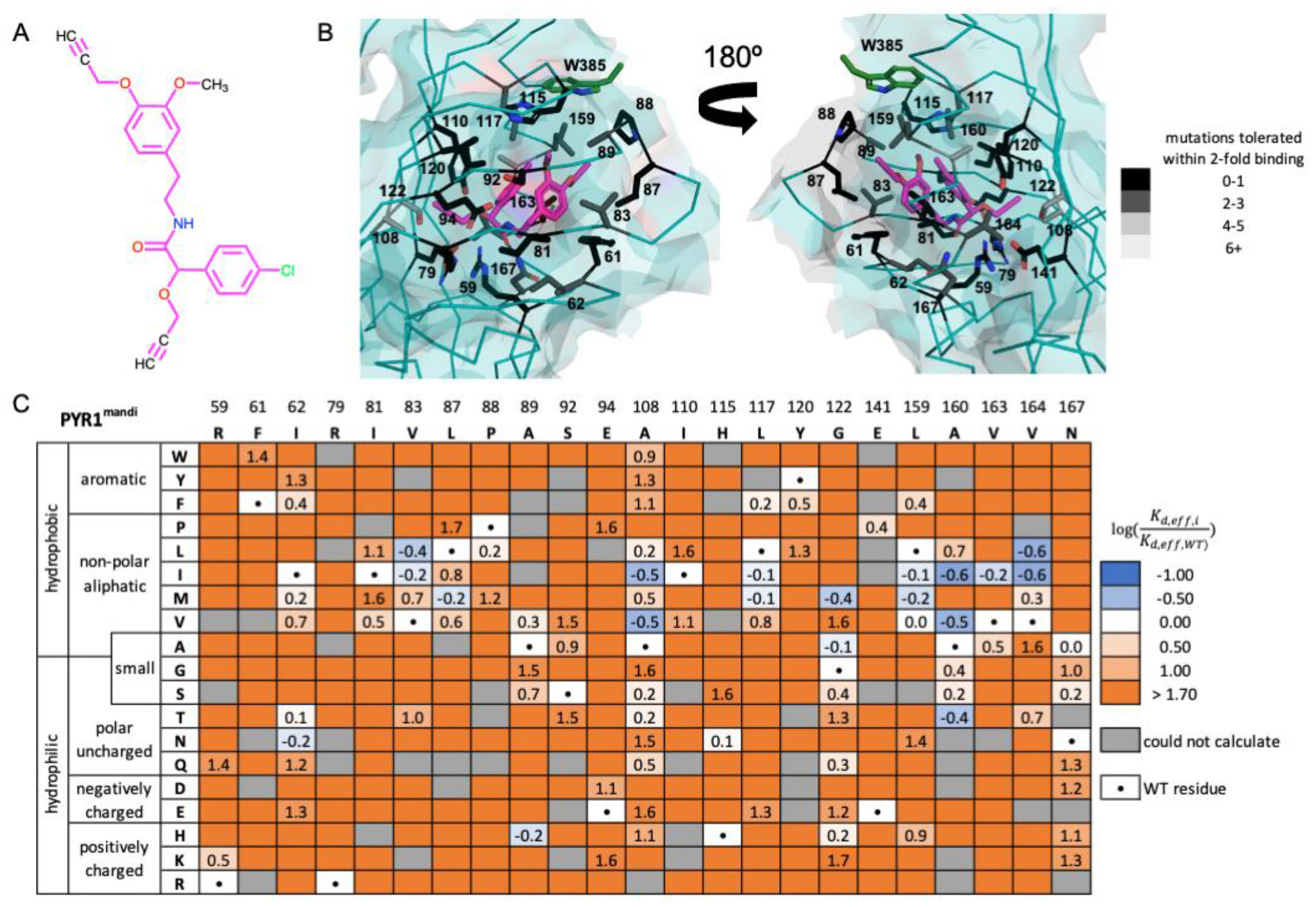
Deep mutational scanning of the mandipropamid engineered biosensor. A. Chemical structure of mandipropamid. B. The mandipropamid ligand binding pocket (PDB 4WVO) color-coded by conservation of position. The side chains of the 23 amino acid positions mutated are shown. C. Heatmap of calculated relative binding affinities of single-point mutations to the mandipropamid sensor, expressed as log(Kd,eff,i/Kd,eff,WT), with blue binding more favorably and orange less favorably.

**Figure 4.**
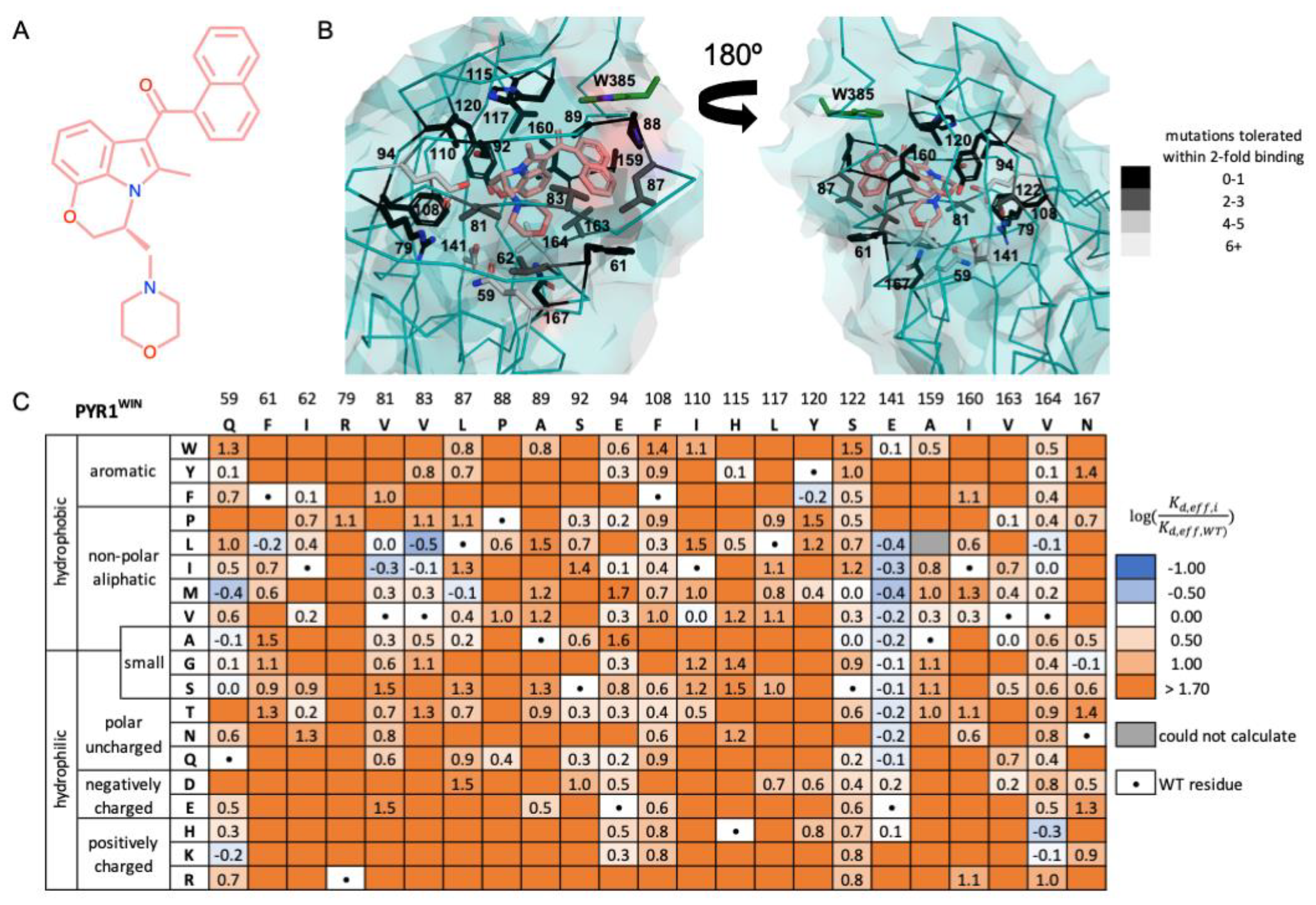
Deep mutational scanning of the WIN engineered biosensor. A. Chemical structure of WIN 55,212-2 ligand. B. WIN sensor ligand binding pocket in the PYL2^WIN^ structure (PDB 7MWN ^13^). Highlighted positions are labeled with the corresponding PYR1 number scheme for consistency. C. Heatmap of calculated relative binding affinities of single-point mutations to the WIN sensor, expressed as log(Kd,eff,iKd,eff,WT). with blue binding more favorably and orange less favorably.

### Sequence-function basis for ligand recognition in the engineered mandipropamid sensor

The agrochemical fungicide mandipropamid is a 23-carbon structure containing two phenyl rings, three ether linkages, and two alkynes (**Figure 3A**). To accommodate such a large molecule, the crystal structure of the engineered PYR1^mandi^ sensor (PDB 4WVO ^2^) shows the ligand compressed in a closed, ‘clamshell’ conformation, with both ring structures pointed up in the pocket (close to HAB1^W385^) and a carbonyl group pointed down (away from HAB1^W385^) (**Figure 3B**). **Figure 3C** contains a heatmap summarizing the results of two replicate deep mutational scanning experiments, considering the 23 ligand-facing residues included in the mutational library. Overall, 77% (319/414) of single point mutations are destabilizing by greater than 50-fold (**Figure 3C**). 9% (34/414) of single-point mutations either improve effective binding affinity or are tolerated within less than a 2-fold reduction in binding affinity (log(Kd,eff,i/Kd,eff,WT) < 0.30). Data from the remaining 61 positions could not be fit to curves to calculate binding affinity.

The PYR1^mandi^ heatmap highlights several key amino acid interactions that are critical to ligand binding. Relative to WT PYR1, the PYR1^mandi^ construct analyzed by deep mutational scanning contains six mutations (Y58H, K59R, V81I, F108A, S122G, F159L) found through directed evolution to enable mandipropamid binding ^2^ (**Table S1**). The reversion mutations (R59K, I81V, A108F, G122S, L159F) are deleterious yet tolerated within a 2.6 to 11-fold reduction in binding affinity. Position R59 is notably one of several positions to tolerate very few other mutations away from wild-type. Other largely conserved positions include P88 and H115, which coordinate a bound water molecule forming the ‘gate’ ^13,32^, as well as R79, E94, and E141 that form intrapocket salt bridges (R79-E94; R59-E141) as evident in the crystal structure of the PYR1^mandi^ complex.

In contrast, other positions accommodate significant mutational flexibility. These positions tend to be near the top of the binding pocket, encoding aliphatic residues such as I81, V83, L87, A108, L117, L159, A160, and V163 that can accommodate mutations to other similarly-sized or slightly larger aliphatics (**Figure 3C**). Position A108 notably can accommodate all but proline, aspartic acid, lysine, and arginine. Nine mutations improved binding affinity by 2.5-fold or greater, corresponding to log(Kd,eff,i/Kd,eff,WT) < -0.40. These were V83L, A108I/V, A160I/V/T, and V164I/L, as well as G122M. All nine mutations were from a smaller residue to a larger, usually hydrophobic residue, suggesting that these substitutions provided additional van der Waals packing and improved shape complementarity to the bound conformer of the mandipropamid ligand.

### PYR1^WIN^ sensor ligand binding

We also performed deep mutational scanning on PYR1^WIN^, an engineered biosensor that recognizes the synthetic cannabinoid (+)-WIN55,212-2 (**Figure 4A**). PYR1^WIN^ was identified through a mutational library screen and contains mutations K59Q, F159A, and A160I away from the PYR1 sequence ^13^. The solved structure of an engineered biosensor bound to WIN55,212-2 shows superficially similar mechanisms of ligand binding with the PYR1^mandi^ structure. The naphthalene and morpholine rings of WIN55,212-2 are folded in a compressed, ‘clamshell’ orientation (**Figure 4B**). Like PYR1^mandi^, the PYR1^WIN^ sensor binds via a water-mediated bond with a key hydrogen bond acceptor in the gate-latch-lock mechanism ^13^. Unlike the PYR1^mandi^ sensor, there is a considerable void at the ‘bottom’ of the pocket at, and adjacent to, position 59.

**Figure 4C** shows a heatmap of deep mutational scanning results for the PYR1^WIN^ sensor for 23 ligand-facing positions. PYR1^WIN^ sequence-binding maps exhibited considerably more sequence permissibility compared to the PYR1^mandi^ sensor. While P88 and H115 are largely conserved, only 50% (205/414) of all single point mutations are destabilizing by greater than 50-fold, and 13% (53/414) either show improved binding or have less than a 2-fold reduction in binding affinity. Unlike the PYR1^mandi^ sensor which tolerated only 59K/R & 141E/P, position 59 in PYR1^WIN^ tolerates the chemically diverse mutations to Q59Y and Q59G with less than a 2-fold reduction in binding affinity, while Q59M, Q59A, Q59S, and the reversion mutant Q59K all improve binding affinity (**Figure 4C**). At position 141, all possible mutations other than to the large or positively charged residues Y,F,P,K, and R are tolerated within a 2-fold affinity reduction, and many mutations improve binding affinity. Since the original K59Q mutation disrupted the K59-E141 salt bridge, the combined mutational results show the relative unimportance of this local electrostatic environment on WIN ligand recognition. This is consistent with the solved structural complex, as E141 does not appear to contribute to hydrogen bonding or charge satisfaction for the WIN ligand. The R79-E94 salt bridge was also disrupted, with position 94 tolerating mutations to W,Y,P,I,V,G,T,Q,D,H, and K within a 4-fold reduction in binding affinity, while R79 remains conserved (**Figure 4C**).

While the electrostatic networks are very different, the PYR1^WIN^ sensor accommodates similar mutational flexibility to the PYR1^mandi^ sensor at the top of the binding pocket, particularly positions V81, V83, V87, and V163 (**Figure 4C**). Mutations V81I, V83L, V83I, and L87M improve effective binding affinity, suggesting that subtle changes in van der Waals packing from aliphatic substitutions can improve shape complementarity. Notably, positions 159 and 160 also near the top of the pocket did not show similar mutational tolerance. The reversion mutations A159F and I160A result in greater than 50-fold worse binding, and at both positions only mutations to valine are tolerated within a 2-fold decrease in binding affinity (**Figure 4C**). The relatively small or large size of the aliphatic residues at those positions is critical, as the naphthalene ring of WIN55,212-2 protrudes close to the alanine at position 159 while space is opened near position 160.

A key difference between the PYR1^mandi^ and PYR1^WIN^ sensors is the relative packing at the bottom of the binding pocket, with the compressed “clamshell” orientation of WIN55,212-2 sitting higher in the pocket and not extending functional groups down like the mandipropamid ligand (**Figure 4B**). Positions S122 and V164 tolerate significantly more variability in the PYR1^WIN^ sensor than in PYR1^mandi^, with both positions tolerating mutation within 50-fold destabilization to nearly any other amino acid (**Figure 4C**). This could be explained by the fact that both S122 and V164 residues are at the bottom of the pocket and distal from the WIN ligand. At position V164, which is located on the inside face of the central alpha helix, mutations to either the positively charged amino acids histidine or lysine or a mutation to the larger aliphatic residue leucine can improve binding affinity. The positively charged residues H/K likely coordinate the unpaired E141, while a larger aliphatic residue is likely filling void space in the lower pocket otherwise occupied by solvent.

In summary, deep mutational scanning identifies mechanisms of ligand recognition common to both engineered biosensors. Protein shape complementarity to the bound ligand is driven by aliphatic residues at the top of the pocket, with gain of function mutations likely improving van der Waals packing. In both sensors, the preservation of key residues satisfies hydrogen bond acceptors not used in coordinating the latch water molecule, with the arginine at position 79 notably conserved across both sensors. In contrast, the other charged residues at positions 59, 94, and 141 had varying differences in conservation between sensors. The PYR1^WIN^ sensor was much more tolerant to mutation, likely owing to the relative paucity of ligand contacts at the bottom of the binding pocket.

However, we were struck by several additional questions raised by the deep mutational scanning results that, if addressed, could improve our structural and physicochemical understanding necessary for *de novo* design of new sensors. First, whether residues not involved in the gate-latch-lock mechanism also formed water-mediated binding interactions that influenced the observed mutational profiles, as some differences in amino acid identities at each position could be rationalized by bound waters. Second, while the electrostatic network surrounding mandipropamid in the PYR1^mandi^ sensor does not tolerate single mutations that disrupt only one end of a salt bridge, it is unclear whether the whole electrostatic network is required for enforcing binding affinity. Third, our deep mutational scanning method analyzed single mutations in isolation, raising questions about whether mutations acted on the same ligand internal conformation and rigid body orientation in the PYR1 binding pocket. If the ligand conformation and orientation was fixed for high affinity sensors, then individual point mutations would be expected to contribute additively to effective affinity gains found during directed evolution, simplifying engineering workflows. Knowledge of likely final fixed conformers could also improve computational design, enabling identification of many additive point mutations during the initial design process.

### Functional role of water networks to ligand recognition in engineered biosensors

The crystal structures of multiple engineered PYR1 sensors, including PYR1^mandi^ and PYR1^WIN^, show the ligand binding via one or two coordinated waters at the mouth of the ligand binding pocket ^13^. To investigate the possible functional role of other coordinated water networks, we performed all-atom MD simulations of the PYR1^mandi^ sensor in complex with HAB1 (see **Methods**). From these MD simulations, we identified both direct and water-mediated non-bonded interactions between protein residues and the ligand using a heavy atom distance threshold of 4Å. We then quantified the relative occupancy R of direct versus water-mediated interactions using the formula R = D-WD+W×I where D is the occupancy of direct non-bonded interactions between a given residue and the ligand, W is the occupancy of water mediated non-bonded interactions between the same residue and ligand, and I is the total occupancy of either direct or water mediated non-bonded interactions. We computed the relative occupancy of these interactions for all residues in both the receptor and HAB1. We considered residues with R < -0.7 to have “dominant” water-mediated interactions (**Figure 5A–B, S2**).

**Figure 5.**
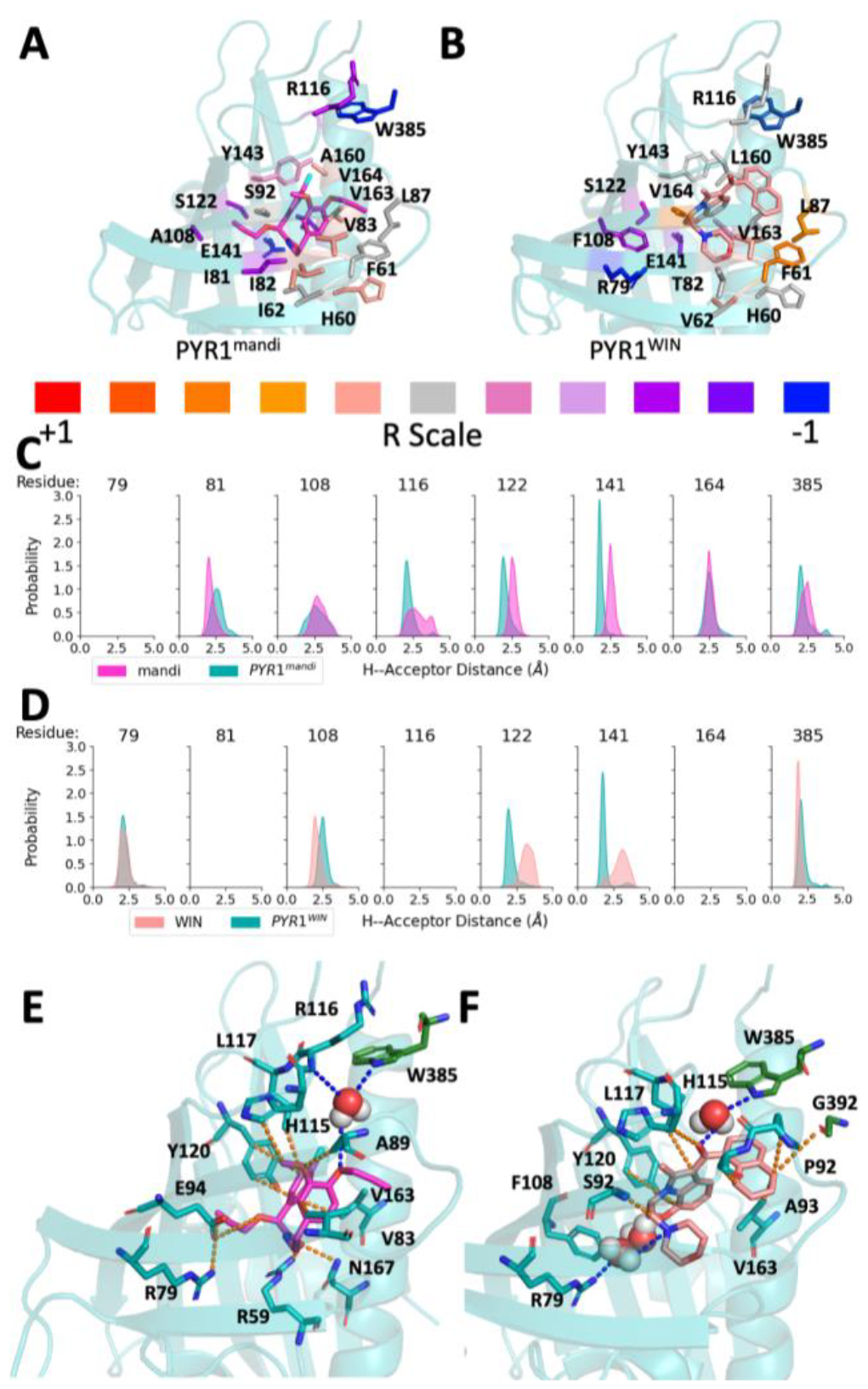
Conserved water coordination at mouth of PYR1^mandi^ and PYR1^WIN^ sensors while other coordinated waters are sensor-specific. A-B. The structures show the relative occupancy of direct vs water-mediated non-bonded interactions between the ligand and both the sensor and HAB1 for PYR1^mandi^ and PYR1^WIN^ sensors. Only residues with |R| > 0.5 are shown (Full data **Figure S2**). Those residues with water dominated non-bonded interactions are identifiable with a deeper blue hue and an R score closer to -1. C-D. Water-mediated H-bonds between the ligand and W385 are stable for both PYR1^mandi^ and PYR1^WIN^ as demonstrated by sharp peaks with a mean of less than 2.5A in the distribution of hydrogen– acceptor distances. Despite several bonds close to the threshold, no other waters were observed as strong and stable in MD simulations for PYR1^mandi^. PYR1^WIN^ does form additional stable, strong, and dominant H-bonds with PYR1 residues R79 and F108. The structure of the binding pocket for both the PYR1^mandi^ (E) and PYR1^WIN^(F) sensors requires the water mediated interactions(blue) with W385 at the top of the binding pocket due primarily to an absence of direct non-bonded interactions(orange) in that region, but only PYR1^WIN^ has a gap in the bottom of the binding pocket which must be compensated for by additional water-mediated h-bonds.

We then sought to determine which of these water mediated interactions are both strong and maintain stability throughout the simulation, suggesting which water molecules are more likely to significantly contribute to the ligand binding mechanism. We explain the determination of “strong” and “stable” h-bonds in the methods below. For PYR1^mandi^, the only protein residues which had “dominant” (R < -0.7) water-mediated interactions which were classified as both strong and stable were those formed with residue W385 on HAB1 (**Figure 5C,E**) which is consistent with the crystal structure. For PYR1^WIN^, the dominant water-mediated interactions that were both strong and stable were found with residues W385 on HAB1 as well as residues R79 and F108 on the receptor (**Figure 5D,F**). The conservation of the coordinated water molecule participating in the gate-latch-lock mechanism at the top of the binding pocket demonstrates the importance of this interaction. The PYR1^WIN^ exclusive coordinated water molecules near the bottom of the binding pocket are likely a compensation for the lack of direct non-bonded interactions in that region. Likely due to the compact nature of the molecule as well as difference in chemical functionalization, the WIN ligand binds in a similar but distinct region of the binding pocket compared to mandipropamid, leaving a much larger cavity within the pocket (**Figure 5E,F)**. These coordinated waters fill the observed void in the binding pocket and provide crucial stabilizing non-bonded interactions which, as suggested by deep mutational scanning, may also be filled with larger aliphatic residues.

### Electrostatic contributions to ligand recognition probed by molecular dynamics and mutational analysis

To address the precise role of the charged residues, and the two salt bridges observed previously in crystal structures, on ligand recognition, we performed MD simulations of several engineered biosensors. To establish that MD results were consistent with the experimental results from deep mutational scanning, we compared the calculated ΔΔG from relative free energy calculations performed with Hamiltonian replica exchange simulations to *log(Kd,eff,i/Kd,eff,WT)* for several mutations in the PYR1^mandi^ sensor and verified that the same trends were observed in experiment and MD (**Figure S4**). In MD, stable salt-bridges R59-E141 and R79-E94 (**Figure 6A**) in the PYR1^mandi^ were observed both with and without HAB1 present with mean occupancies of 91% and 100% respectively (**Figure S5)**. This suggests that the formation of these H-bonds precede the secondary HAB1 binding event and are potentially necessary for initial PYR1-mandi complex formation. Additionally, R59 forms a stable H-bond with a mean occupancy of 98% with the mandipropamid ligand, appearing to stabilize the binding pocket as demonstrated by the significant reorientation of the ligand in the absence of this H-bond (**Figure 6B, S5**).

**Figure 6.**
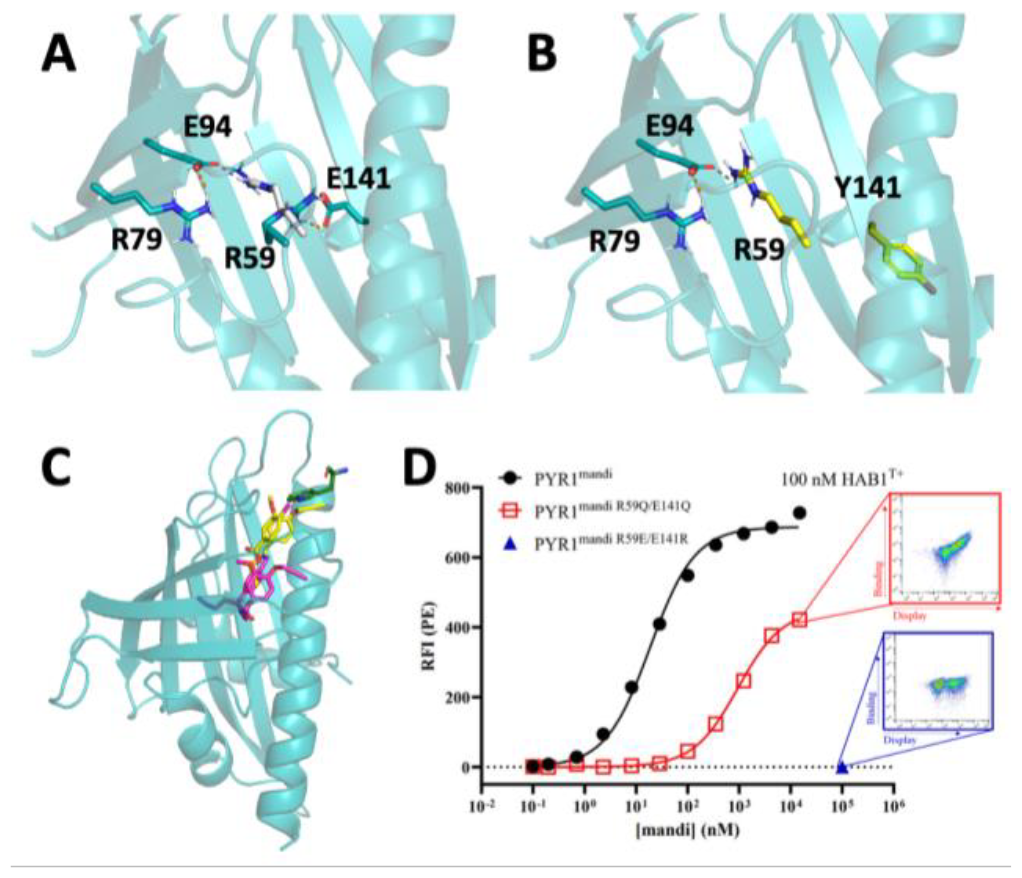
Electrostatic interactions facilitate H-bonding to stabilize the bottom of the mandi ligand binding pocket, which is necessary for HAB1 binding to the receptor-ligand complex. ***A***. There are two primary salt bridges present within the binding pocket of PYR1^mandi^ with ligand bound, R79–E94 and R59–E141 (shown in cyan). In the absence of ligand, R59 (shown in white) is oriented up into the space ligand will occupy, forming a salt bridge with E94. These are observed in MD simulations as well as in the crystal structure (PDB 4WVO) **B**. In MD simulations of the ligand complex with PYR1^mandi/E141Y^, the E141Y mutation prevents salt bridge formation with R59, and R59 orients upward similar to the absence of ligand. Y141 instead adopts an extended, nearly perpendicular conformation (shown in yellow) as its backbone loop shifts. **C**. In complex with PYR1^mandi/E141Y^, the ligand (shown in yellow) exhibits a significantly different binding conformation than the WT PYR1^mandi^ complex. We predict that the absence of an anchoring H-bond between the ligand and the reoriented R59 in the PYR1^mandi/E141Y^ variant is the primary cause for this change in ligand orientation. The position of mandipropamid in PYR1^mandi/E141Y^ causes a direct clash with the predicted location of W385 thus suggesting that this mutation would disrupt the gate-latch-lock mechanism and prevent ternary complex formation. **D**. Yeast surface display titrations of PYR1^mandi^, PYR1^mandi R59Q/E141Q^, and PYR1^mandi R59E/E141R^. Relative fluorescence intensity (RFI) is determined by secondary labeling using streptavidin-phycoerythrin after initial labeling with indicated ligand concentration and 100 nM biotinylated HAB1^T+^. Error bars represent 1 s.d. In relative fluorescence for n = 2 replicates. Inset cytograms show binding of HAB1^T+^ versus display of PYR1 variant on the surface of yeast. The cytogram for PYR1^mandi R59Q/E141Q^ shows the positive result of binding and the cytogram of PYR1^mandi R59E/E141R^ shows the lack of binding for that variant.

We performed simulations of the PYR1^mandi^ sensor with the mutation E141Y to investigate how the binding pocket changes in the absence of these stabilizing salt-bridge interactions. PYR1^mandi/E141Y^ was demonstrated to have significantly reduced binding affinity in deep mutational scanning (**Figure 3C**), presumably by breaking the R59-E141 salt bridge. In simulations of PYR1^mandi/E141Y^, the occupancy of the R59-Y141 H-bond is less than 10%, compared to mean occupancy of 91% for the R59-E141 H-bond in PYR1^mandi^ (**Figure S5**). When E141 is mutated to tyrosine, R59 adopts a conformation like that of the apo state (**Figure 6A-B**) which prompts the formation of the R59-E94 H-bond, which is also present in Apo PYR1^mandi^ (**Figure S5**). These differences in the H-bond network within the binding pocket prevent the formation of the H-bond between the ligand and R59. The absence of this H-bond results in the destabilization of the ligand in the WT binding conformation and subsequent movement of the ligand to the top of the binding pocket (**Figure 6B and S5-S6**). Though the PYR1^mandi/E141Y^ variant results in minimal reorientation of the backbone of PYR1, this ligand binding location is not conducive to the formation of the PYR1-HAB1 complex (**Figure 6A, S7**). This suggests that R59 serves as an anchor for mandipropamid, ensuring binding occurs in an orientation which allows for the subsequent binding of HAB1 to the complex. Overall, molecular dynamics simulations support that the formation of the R59-E141 salt bridge via electrostatic interaction enables the H-bonding of R59 to mandipropamid necessary for ligand recognition by correctly positioning residue R59 for binding.

To build on the mechanistic insights of single mutations gained from simulation, we experimentally probed double mutants of the R59-E141 and R79-E94 salt bridges. While single mutations to R59 or E141 in PYR1^mandi^ leave one half of an unsatisfied salt bridge intact, simultaneous mutation at both ends can explore the full impact of the electrostatic interaction. We hypothesize that if the effect of the salt bridge is simply the maintenance of electrostatic neutrality in the pocket, then swapping the salt bridge to R59E/E141R should still maintain binding. If, on the other hand, R59 contributes a critical H-bond donor to the ligand, this could be partially satisfied by a residue like Gln containing an H-bond donor. To test these hypotheses, we constructed the double mutants PYR1^mandi/R59E/E141R^ and PYR1^mandi/R59Q/E141Q^ and performed titrations of different concentrations of mandipropamid (**Figure 6D, S8**). The mutation E141Q was chosen as a non-charged isostere maintaining charge neutrality. In attempting to swap the charge across the salt bridge with mutations R59E and E141R, we found that binding was ablated, even at 100 μM concentration of mandipropamid. Removing both charges with simultaneous glutamine mutations resulted in functional binding, albeit with a significantly worse K_D,eff_. These results show that maintenance of electrostatic neutrality alone is not enough to enable binding. However, electrostatic neutrality combined with the presence of a hydrogen bond donor can maintain mandipropamid recognition. These results agree with our molecular dynamics simulations, in that it is the H-bonding capability at position R59 that is essential for ligand recognition.

The R79-E94 salt bridge provides both electrostatic neutrality and potential PYR1 local secondary structure stabilization. MD simulations reveal that R79 forms a h-bond with the main chain carbonyl of F52 with >90% occupancy in both apo and ligand bound PYR1^mandi^ (**Figure S9**), suggesting a potential structural role for this residue. This is consistent with the deep mutational scanning results that residue R79 is critical in both sensors. To test this hypothesis, we prepared a series of double mutants that maintain electrostatic neutrality. If the PYR1 structure is not impacted, these mutants should still result in functional binders, as neither residue appears to be hydrogen bonding with the ligand in PYR1^mandi^ or PYR1^WIN^. However, all double mutants tested (PYR1^mandi/R79E/E94R^, PYR1^mandi/R79A/E94A^, PYR1^mandi/R79T/E94Q^, PYR1^WIN/R79T/E94Q^, PYR1^WIN/R79I/E94L^) resulted in a total loss of ligand binding (**Figure S10**). To determine whether mutation at R79 results in a PYR1 not able to maintain HAB1 recognition, we tested binding of PYR1 mutants in the presence of biotinylated HAB1^T+^ and in the absence of ligand. The parental PYR1^mandi^ and PYR1^WIN^ sensors, and all R79 mutants assayed, bound weakly at low micromolar HAB1^T+^ concentrations and with no difference compared to the parental sensors (**Table S4**). Combined, these results suggest that R79 is important for maintenance of local, though not global, structure necessary for ligand binding.

### Conformer selection of high affinity sensors

While a solved crystal structure captures a single ligand conformation in the binding pocket, a protein-ligand complex could accommodate different modes of binding to different ligand conformers. To address this, we used MD simulation to sample the mandipropamid and WIN55,212-2 ligand conformations adopted in solution and compared these to the conformers sampled in complex with their engineered PYR1 variant. During 3 independent 300 ns MD simulations of PYR1^mandi^ and PYR1^WIN^ in complex with the ligand and HAB1 only one ligand conformer was observed which matches in the published crystal structures 4WVO (**Figure 7A)** and 7MWN (**Figure 7B**), respectively. Using temperature replica exchange MD simulation we sampled conformations of both ligands in solvent allowing us to rank the energy clustered ligand conformers (See **methods** for additional details). We found that the observed conformer is among the most energetically favorable solution conformation (**Fig 7C-D**). The free energy penalty relative to the lowest energy conformer in solution is below 1 K_B_T for both ligands, and both are estimated roughly to pay less than 1 kcal/mol free energy penalty to go from all conformers in solution to the complexed conformer ^39^. While the low-energy protein-ligand complex does not necessarily bind the lowest-energy ligand conformer in solution, the conformations which are selected are not unduly strained or otherwise highly unlikely to occur in solution.

**Figure 7:**
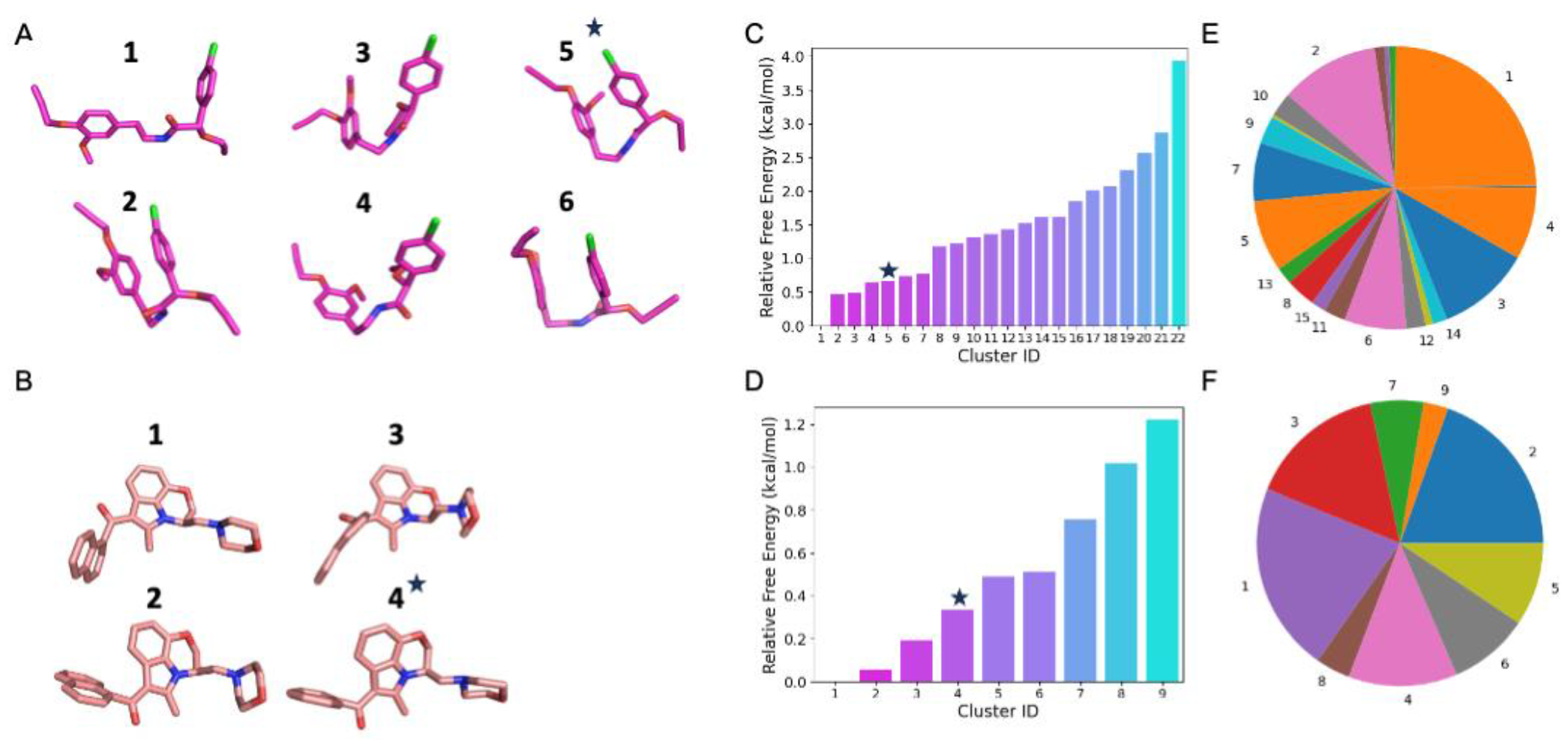
PYR1^mandi^ and PYR1^WIN^ select for a single low-energy ligand conformer. We evaluated the conformational flexibility of mandipropamid (A) and WIN55,212-2 (B) in solvent using temperature replica exchange molecular dynamics which allowed us to rank clusters of conformers for each ligand in solution. We hypothesize that mandipropamid conformer cluster 11 could accommodate mutations requiring the rotation of a methyl propargyl ether group.The conformer which is sampled when the ligand is in complex with PYR1 is indicated with a star and is the 5th and 4th lowest free energy conformation for mandipropamid (C) and WIN (D) respectively. E-F. Probability of observed conformer cluster in the 300 K replicate during temperature replica exchange molecular dynamics in solvent for mandipropamid and WIN55,212-2).

Many protein-ligand complexes gain increasing ligand binding affinity through the accumulation of additional mutations that each individually improve binding affinity ^40,41^. A first-order approximation for additivity of mutational effect is that mutations do not change ligand conformer nor rigid body orientation. Thus, if PYR1^mandi^ and PYR1^WIN^ each bind only a single ligand conformer in a constrained rigid body orientation, then further mutations should be additive unless part of the conformer can rotate.

To probe whether the additional mutations would continue to select for a single conformer, we performed experiments on multi-mutants of both sensors to compare the predictive benefit of additive mutations to experimentally-determined changes in binding affinity. The mutant of PYR1^WIN^ had the mutations Q59M/Y120F/E141M (**Figure 8A left**), all of which deep mutational scanning indicated improved binding compared to WT PYR1^WIN^. Ligand titrations at a constant HAB1^T+^ concentration were performed for this mutant and the binding affinity was compared to the original sensor, as in Figure 3C and 4C (**Figure 8B left**). PYR1^WIN Q59M/Y120F/E141M^ had a K_d,eff_ that was ∼10-fold lower than the original PYR1^WIN^ sensor, showing this expected additive effect and suggesting that WIN was in the same conformation for each individual mutation (**Figure 8C left**).

**Figure 8:**
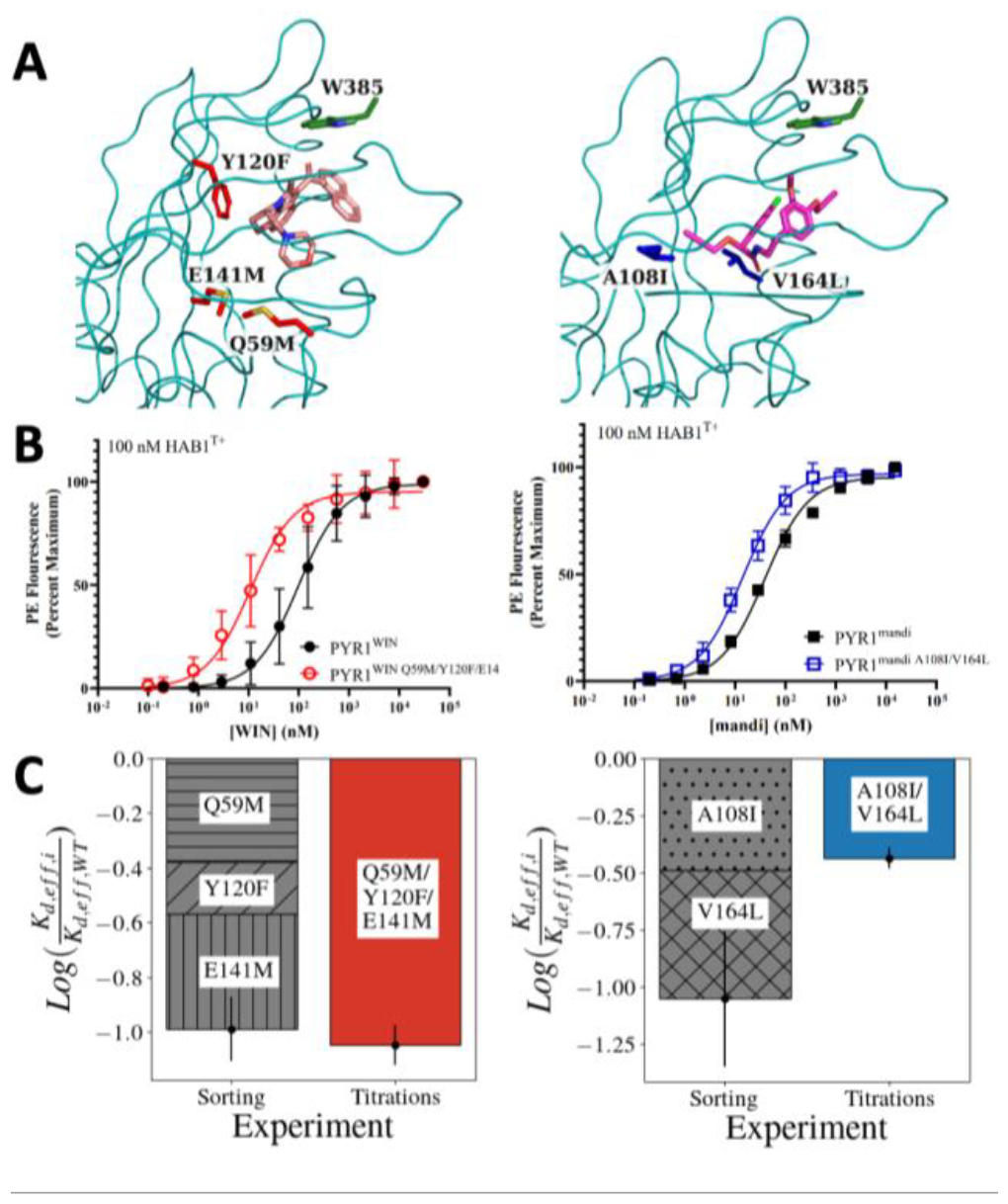
Degree of mutational additivity indicates conformer selection in engineered biosensors. A) Structures of PYR1^WIN Q59M/Y120F/E141M^ (left) and PYR1^mandi A108I/V164L^ (right). The PYR1 backbone is shown as a teal ribbon. Ligands and mutated residues are shown as sticks. B) Yeast surface display titrations of multi-mutant biosensors for WIN (left) and mandi (right) compared with original sensor. PE fluorescence is determined by secondary labeling using streptavidin-phycoerythrin after initial labeling with indicated ligand concentration and 100 nM biotinylated HAB1^T+^. Error bars represent 1 s.d. in relative PE fluorescence for n=4 (two technical replicates for two biological replicates). C) Predicted additive improvement of mutations in binding affinity compared to experimentally observed relative binding affinity. PYR1^WIN Q59M/Y120F/E141M^ (left) is additive while PYR1^mandi A108I/V164L^ (right) is not. Error bars represent the summation of mean absolute error for each mutation (gray) or mean absolute error for titrations (colored).

For the mandi sensor, titrations of the double mutant A108T/L159F matched the decrease in binding affinity compared to WT PYR1^mandi^ predicted by deep mutational scanning (**Figure S11 left**). In contrast, PYR1^mandi A108I/V164L^ and PYR1^mandi G122M/A160I^ were expected to increase binding affinity compared to PYR1^mandi^. However, titrations of PYR1^mandi A108I/V164L^ (**Figure 8A right**) indicate that the double mutant does decrease K_d,eff_, but by a different magnitude than would be predicted for fully additive mutations (**Figure 8B,C right**). Similar results were observed for PYR1^mandi G122M/A160I^ (**Figure S11 right**). In these double mutants without additivity, the combined mutations were close to each other in space and present in the pocket where the mandipropamid molecule has a freely rotatable methyl propargyl ether group sticking into the bottom of the pocket, suggesting that each mutation acts on a different, or constellation of, ligand conformation(s) in the same approximate rigid body orientation (**Figure 7A**).

### Computational design of WIN binding from a fixed selected conformer

If high-affinity protein-ligand binders select for a single ligand conformer, then correctly identifying likely ligand conformations and placements in the binding pocket would be crucial for design success. To test whether computational design could result in functional PYR1 biosensors when provided a fixed ligand conformer and rigid body orientation, we performed Rosetta FastDesign ^42^ on the experimentally determined PYL2-WIN55212,2 crystal structure using the PYR1^WIN^ mutations reverted to wild-type PYR1/PYL2 sequences. FastDesign suggested the mutations S92V (deleterious in the high affinity sensor; **Fig 4C**) and F159AMVI (allowable; **Fig 4C**). We tested combinations of mutations at these positions along with the known electrostatic-altering K59Q mutation observed in the PYR1^WIN^ sensor. All six designs bound WIN using our yeast display assay (**Fig 9, S12)**, and two sensors with an F159A mutation bound with a high nM limit of detection (**Fig 9**). Overall, these results show that physically based energy functions (Rosetta) are sufficient for identifying binding sequences if a compatible ligand conformation and rigid body orientation can be identified.

**Figure 9:**
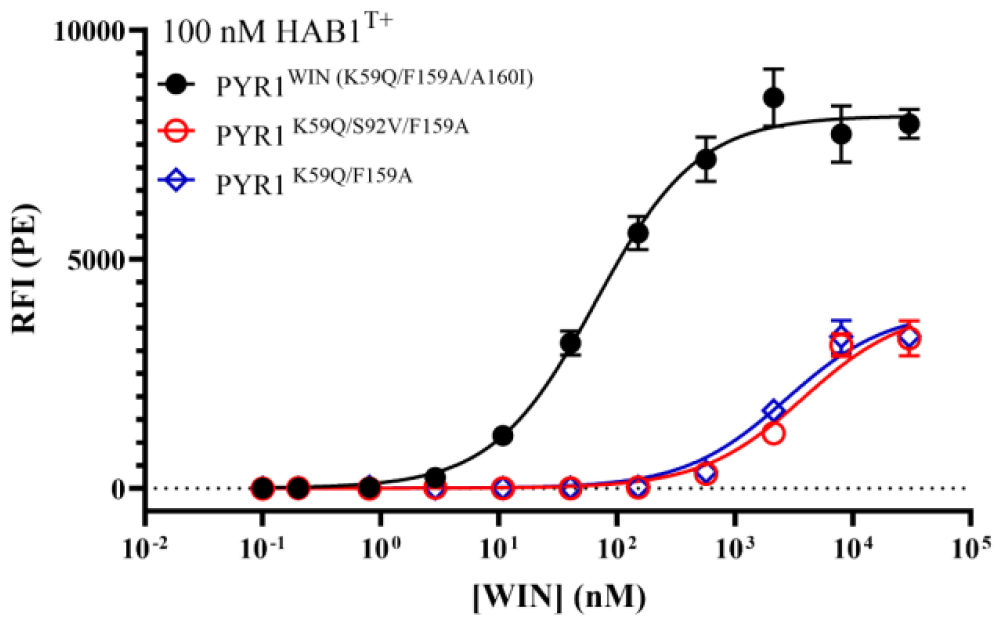
Computational design of PYR1-ligand binding from a selected fixed conformer. Yeast surface display titrations of computationally designed PYR1 biosensors shown in red and blue binding WIN55,212-2. The wild-type PYR1^WIN^ sensor shown in black. PE fluorescence is determined by secondary labeling using streptavidin-phycoerythrin after initial labeling with indicated ligand concentration and 100 nM biotinylated HAB1^T+^. Error bars represent 1 s.d. in relative PE fluorescence for n=2 (two technical replicates for one biological replicate, second biological replicate in figure S12).

## Discussion

In this work we analyzed the structural, sequence, and mechanistic determinants of binding in two engineered PYR1 biosensors. We performed deep mutational scanning, used MD simulation to explore conformer selection and mechanisms of ligand binding, generated multi-mutants to test distinct biophysical hypotheses, and used computational design to apply these insights to design new PYR1^WIN^ sensors.

Some lessons for computational design will apply to engineering PYR1 for binding to other novel ligands, and may apply to computational design of protein-ligand binding more broadly. In the PYR1 scaffold, mutations at the top of the binding pocket to different sized aliphatics appear to increase shape complementarity for improved binding affinity. Additionally, hydrogen bonds with available ligand functional groups can stabilize binding, particularly at position 59 to anchor the ligand in the bottom of the binding pocket. The nature of the H-bond networks between ligand and protein are closely related to the electrostatic environment in the binding pocket, as the charge of residues forming hydrogen bonds must be balanced across the pocket by salt bridges. Balancing charge in the pocket is important for high-affinity binding, but not sufficient alone if ligand-stabilizing H-bonds are removed. While we determined through MD that water-mediated latch closure involving HAB1^W385^ is essential to ligand binding in engineered PYR1-HAB1 complexes, additional water molecules can play a role in filling the bottom of the binding pocket for compact ligands, such as bound WIN55,212-2, forming stable water-mediated interactions between protein and ligand. These lessons suggest that to engineer PYR1 binding to novel ligands one should focus on satisfying ligand H-bond acceptors, balancing charge across the binding pocket, and targeting mutations to complement ligand size and shape to the top of binding pocket. Indeed, following these lessons led to computational design of sequence-distinct PYR1 proteins which recognize WIN55,212-2 with high nanomolar limit of detection on the yeast surface.

High affinity biosensors are likely evolved for a single ligand conformer, or closely-related cluster, in the same rigid body orientation through the accumulation of additive beneficial mutations. This hypothesis is also supported by our MD simulations observing that PYR1^mandi^ and PYR1^WIN^ each bind a single ligand conformer. This idea of conformer selection by the accumulation of beneficial additive mutations can be seen in the development of the PYR1^mandi^ sensor ^2^ and the broader screening of organophospates and cannabinoids ^13^. These papers showed that the PYR1 binding pocket is unusually pliable, with medium affinity binders able to be identified from a small number of mutations. In Park et al, an early PYR1 sensor binding mandipropamid contained only the mutations K59R, S122G, and F108A and bound the ligand with low micromolar responsiveness, while additional affinity maturation of the sensor added the mutations Y58H, V81I, and F159L ^2^. However, the deep mutational scanning data from this work indicates that position 108 of PYR1^mandi^ can tolerate significant mutation, and is improved by substitution to the slightly larger non-polar aliphatic residues isoleucine and valine.

Under the mechanism that acquiring more additive mutations fixes the ligand to only a handful of similar ligand conformations, we hypothesize that the original F108A mutation opened the ligand binding pocket to accept the mandipropamid molecule, and then additional mutations selected for a narrower range of allowed conformers. This can also be understood as a tradeoff between enthalpy and entropy in the overall free energy of the system, in which restricting the allowed ligand conformers increases the free energy by decreasing the entropy, but is outweighed by the more negative enthalpy from improved binding affinity. If the accumulation of additive mutations fixes conformer selection in the binding pocket, then sensors with fewer mutations are more likely to bind their ligand weakly and less specifically, and sensor variants containing more mutations would be easier to further mature through directed evolution because each additional mutation is more likely to be additive.

Another crucial computational design consideration is the rigid body placement of the appropriate conformer with respect to the protein binding pocket. The mutational and MD studies here reiterated the known importance of the bound water coordinated by Trp^385 13,35^. Because for all known biosensors a ligand H-bond acceptor coordinates this bound water, the design process for PYR1 is considerably simplified compared to scaffolds without an appropriate anchor point. Indeed, aligning the known WIN conformer to its known H-bond acceptor geometry, followed by Rosetta FastDesign ^42^, is sufficient to generate designs that recognize WIN with a nanomolar limit of detection. Developing new deep learning methods to learn and design small molecule-protein interactions is in vogue ^15,43–48^. We suggest that comparable attention should be placed with the choice of ligand conformer and rigid body orientation.

## Conclusion

In this work we analyzed two engineered biosensors to understand how the same PYR1 protein scaffold can be mutated to bind two ligands with very different structural features. We evaluated the sequence determinants of binding using deep mutational scanning and performed molecular dynamics analysis to elucidate key mechanisms of binding for multiple ligands. This analysis provides insight into how electrostatic networks complement different ligand structural features, how the directed evolution of protein-ligand interactions can promote selection of a specific conformer, and how proper sampling of plausible conformers is critical for successful computational design. While the insights from this work can directly inform PYR1 scaffold engineering for novel ligand biosensors, selection of a limited conformer repertoire is likely a trait of many high affinity protein-ligand interactions and can be applied generally for computational design.

## Methods

### Construction of PYR1^mandi^ and PYR1^WIN^ mutational libraries

Single-site saturation mutagenesis libraries of the PYR1^mandi^ and PYR1^WIN^ sensors were created using comprehensive nicking mutagenesis, in which each specified position was mutated to every other amino acid plus stop codon^49^. For the PYR1^mandi^ sensor, plasmid pJS624 was mutated using NNK primers at positions 59, 60, 61, 62, 79, 81, 83, 87, 88, 89, 91, 92, 94, 108, 109, 110, 115, 117, 120, 122, 141, 158, 159, 160, 163, 164, and 167 to create library L054. For the PYR1^WIN^ sensor, plasmid pRMC005 was generated by Golden Gate assembly of synthetic dsDNA (eBlock, Integrated DNA Technologies) into vector pND003 (Daffern et. al. 2023) and mutated using NNK primers at positions 59, 61, 62, 79, 81, 83, 87, 88, 89, 91, 92, 94, 108, 109, 110, 115, 116, 117, 120, 122, 141, 158, 159, 160, 163, 164, 167 to create library L058. Primer sequences are listed in Table S2. Library plasmids were transformed into chemically competent EBY100 cells as described (Medina-Cucurella and Whitehead 2018) and stored as 1 ml stocks at OD_600_=1 in yeast storage buffer at -80°C.

L054 replicates A and B were made as two separate library reactions, then screened on different days. L058 was a single library where two replicates were screened on different days.

### Preparation of HAB1 for binding assays

An N-terminal truncation of HAB1 (ΔN-HAB1) purified and stored in saturated ammonium sulfate at 100μM, as described in Steiner et al^50^. For use in yeast surface display assays, the protein was centrifuged at 17,000 x g for 10 minutes to pellet. The supernatant was removed by pipette and discarded, and the pellet was resuspended an equivalent volume of ice-cold CBSF++ (CSBF: 20mM sodium citrate, 147mM NaCl, 4.5mM KCl, 0.1% w/v bovine serum albumin, pH 8.0 adjusted with 1M sodium hydroxide, sterile filtered), 1mM freshly dissolved DTT, 1mM TCEP, pH 8.0). The resuspended HAB1 was then desalted using a Zeba™ spin desalting column (Thermo) equilibrated with CBSF++ and stored on ice.

### Yeast surface display of PYR1 variant libraries

Yeast surface display of PYR1 variant libraries were performed according to Steiner et al., with the following modifications^50^. 1ml yeast stocks of libraries L054 and L058 were thawed, centrifuged at 16,000 x g for 1 min, resuspended in 1ml of SDCAA plus 10 μl Pen-Step (10,000 U/ml, Life Technologies), and grown for 4-6h at 30°C with shaking at 300rpm. Expression was induced by resuspending the SDCAA culture in 1 ml SDGCAA (1 part SDCAA to 9 parts SGCAA) plus 10 μl Pen-Step to OD_600_=1 and growing for 20-22h at 22°C, after which cells were resuspended in 1ml CBSF (20mM sodium citrate, 147 mM NaCl, 4.5mM KCl, 1 g/L bovine serum albumin) at OD_600_=2.

For each ligand labeling concentration, 200 ul cells at OD_600_=2 were mixed with 20 μl ligand diluted in DMSO and 100 μl prepared HAB1 diluted in CSBF++, to a final volume of 1ml in CBSF buffer, ensuring a consistent 1:50 ligand stock dilution and 1:10 HAB1 stock dilution across all reactions. Reactions were incubated at room temperature on a benchtop plate agitator for 30 minutes. Reactions were then centrifuged at 16,000 x g for 1 min to pellet, cells were washed with 1ml CBSF, and centrifuged again. After the supernatant was removed by pipet, reactions were labeled with 12μl anti-c-*myc*-FITC (Miltenyi Biotec), 5μl SAPE (streptavidin-R-phycoerythrin, Life Technologies), and 378 μl CBSF. Labeling reactions were incubated for 10 minutes on ice protected from light, then centrifuged and washed with 1ml CBSF as above, then centrifuged and stored on ice with the supernatant removed.

Cell sorts were performed on a Sony SH800S cell sorter (Sony Biotechnology), with cell pellets resuspended in 1ml of CSBF immediately before reading. For each sample, roughly the top 25% of displaying cells by binding signal were collected. Cell sorter parameters and full sorting statistics are listed for each sensor replicate in the supplementary data file PYR1_DMS_supplemental_data&primers.xlsx. Collected cells were suspended in 5ml SDCAA media plus 50 μl Pen-Step and incubated at 30°C with shaking at 300 rpm for 30–40 hrs, before freezing as 1 ml cell stocks at OD_600_=4 in yeast storage buffer at -80°C.

### Deep sequencing preparation

Cells samples collected from library sorting were prepared for deep sequencing as described in Medina-Cucurella and Whitehead (Medina-Cucurella and Whitehead 2018), using a Zymo Yeast Plasmid Miniprep II kit (Zymo Research) and a Monarch PCR & DNA Cleanup kit (NEB) with the following changes. Samples were amplified using inner primers ACL-P1060 and ACL-P1061 in a 40μl PCR reaction using Q5 Hot Start 2x Master Mix (New England Biolabs) for 20-25 cycles at an annealing temperature of 64°C. 5 μL of PCR product from the inner primer amplification was cleaned using 5μL Exonuclease I (NEB) and 2μL rSAP (NEB), incubating for 15 min at 37°C then 15 min at 80°C. 1.6μL of cleanup product DNA was carried forward to the 2^nd^ PCR reaction using Illumina TruSeq small RNA adapters in a 25μl Q5 reaction for 20-25 cycles at an annealing temperature of 64°C. Samples were purified using Agencourt Ampure XP beads (Beckman Coulter), quantified using PicoGreen (ThermoFisher), pooled, and sequenced on an Illumina MiSeq using 2 × 250 bp paired-end reads by the Rush University Medical Center sequencing facility.

### Maximum likelihood estimation analysis of sequencing data

Variant read counts obtained from deep sequencing were processed by a maximum likelihood estimation method as in Petersen et. al.^38^.

### Construction and yeast surface display of isogenic variants

Individual PYR1 sequence variants for isogenic titrations, analysis of electrostatic variants, and conformer selection experiments were constructed by Golden Gate assembly of an eblock DNA sequence (Integrated DNA Technologies) into vector pND003 (Daffern et. al. 2023). All plasmids are listed in table S3.

Preparation of cell cultures for yeast surface display titrations of isogenic variants was performed as above for the display of PYR1 libraries. Yeast surface display titrations were performed according to Steiner et al AiChE 2020, with the following modifications. For each ligand labeling concentration, 5ul cells at OD_600_=2 were mixed with 1μl ligand diluted in DMSO and 5μl prepared HAB1 diluted in CSBF++, to a final volume of 50μl in CBSF buffer, ensuring a consistent 1:50 ligand stock dilution and 1:10 HAB1 stock dilution across all reactions. Reactions were incubated at room temperature on a benchtop plate agitator for 30 minutes. Reactions were then centrifuged at 2,500 x g for 5 min to pellet, cells were washed with 200μl CBSF, and centrifuged again. After the supernatant was removed by flicking, reactions were labeled with 0.6μl anti-c-*myc*-FITC (Miltenyi Biotec), 0.25μl SAPE (streptavidin-R-phycoerythrin, Life Technologies), and 49.15μl CBSF. Labeling reactions were incubated for 10 minutes on ice protected from light, then centrifuged and washed with 200μl CBSF as above, then centrifuged and stored on ice with the supernatant removed.

Binding measurements were performed on a Sony SH800S cell sorter (Sony Biotechnology), with cell pellets resuspended in 100μl of CSBF immediately before reading. Sample analysis was performed using FlowJo 10, and binding parameters were determined in Graphpad Prism 10.1.0 using the specific binding with Hill slope nonlinear regression function.

### Molecular dynamics simulations

All molecular dynamics (MD) simulations were performed using GROMACS 2020.6. All proteins were parameterized using the amber ff14sb protein force field and all ligands were parameterized using GAFF. The protonation state was determined using the H++ server at a pH of 7. All mutant proteins were prepared using MODELLER. Hydrogen mass repartitioning was applied using ParmED. The simulation box was constructed to maintain a minimum 1 nm distance to the periodic boundary condition, and sodium and chloride ions were added to neutralize the system and maintain 0.15M salt concentration. The procedure for all simulations was as follows: energy minimization to 100 kJ/mol/nm energy threshold, 100 ns NVT equilibration with the berendsen thermostat, and 100 ns NPT equilibration with the berendsen thermostat and barostat. Production simulations were 300 ns, unless otherwise specified, and were run with a 4 fs time step using the Parrinello-Rahman barostat and v-rescale thermostat at 1 atm and 300 K. All input simulation files are provided at the Github repository shirtsgroup/PYR1_Design.

The initial structures for PYR1^Mandi^ and PYR1^WIN^ in complex with HAB1 and the ligand were taken from x-ray crystal structures (PDB 4WVO and 7MWN respectively). We ran simulations of PYR1^Mandi^ and PYR1^WIN^ in the absence of HAB1 as well as apo PYR1 simulations in the absence of both HAB1 and the ligand. The same PYR1 structure was used to initiate all simulations with either HAB1 or both HAB1 and the ligand removed. Mutant simulations used the WT crystal structure as a base and then a mutation was performed using MODELLER. A 5-50 ns equilibration period was included in all simulations to allow for conformational changes to be made from these artificial augmentations to the initial structure as well as to allow the system to settle at the equilibration temperature and pressure. The equilibration period was determined by the lack of systematic change in the protein backbone RMSD.

We analyzed simulations for non-bonded interactions which were broadly defined as a heavy atom distance of less than 4 Å using MDTraj *compute_distances* function. In order to quantify the stability of the water-mediated hydrogen bonds, we computed the distance distribution between the hydrogen and the acceptor in the water mediated H-bond for all residues with dominant water-mediated interactions. We computed the mean distance to quantify the strength of the H-bond. and the standard deviation of the distance distribution to quantify the relative stability of the H-bond, both between the water molecule and the protein residue as well as the water molecule and the ligand. “Strong” H-bonds were classified as having a relatively short mean distance of 2-2.5 Å while weak H-bonds had a longer mean distance of 2.5-4 Å. “Stable” H-bonds were defined as having a standard deviation in bond length of <0.45Å. The numerical ranges for classifying strong and stable H-bonds were set to include 95% of analyzed water– residue H-bonds (**Figure S3**). We included H-bonds with solvent exposed residues as well as residues within the binding pocket which were of the same type as those engaging in water-mediated H-bonds with the ligand.

In addition to standard MD, we also performed alchemical relative free energy (RFE) calculations as well as temperature replica exchange molecular dynamics simulations (TREMD). The crystal structure conformation was used as the WT for RFE simulations and then hybrid topologies and coordinates were generated for mutated residues using PMX. The same energy minimization and equilibration steps were carried out for each of the 18 intermediate λ states. Production simulations were completed using hamiltonian replica exchange using a 2 fs time step and were completed for 25 ns for each replicate. Analysis was completed using the Alchemlyb package with reported ΔΔG estimates computed using MBAR. The TREMD simulations were run with just the ligand in solvent with sodium and chloride ions to maintain a 0.15 M salt concentration. 27 temperature replicates were used varying from 300 K to 450 K with conformational swaps allowed every 2 ps. All conformational analysis came fromonly the 300 K replicate. Sampled ligand conformations were first sorted based on sampled dihedral angles computed using MDTraj and these conformations were then clustered using nearest neighbor clustering on pairwise heavy atom RMSD. The relative free energy of a conformer can be computed since these MD simulations maintain the Boltzmann distribution, thus G -kBTlog(ppref) in which pref is the probability of the most probable conformation and p is the probability of any given conformation. The analysis code can be found at the Github repository shirtsgroup/PYR1_Design.

### Computational design and experimental validation

For computational design, the PYL2^WIN^ structure PDB 7MWN was manually stripped of all accessory ions and water molecules, other than the key water coordinating by the gate-latch-lock binding mechanism, leaving the WIN55,212-2 ligand in place on a separate chain. To eliminate WIN55212-2 binding, we reverted the PYL2^WIN^ sequence to the wild-type PYL2 sequence by mutating Q59K, A165F, and I166V in PyMOL. Computational design using FastRelax ^42^ with the default energy function was performed using PyRosetta4 version 2021.26+release.b308454c455dd04f6824cc8b23e54bbb9be2c dd7, performing design on 9 subtle variations of ligand alignment to the water molecule. Output designs were screened using PyMOL to ensure the ligand formed a polar contact with the water molecule and were scored by Rosetta on interface energy, buried unsatisfied hydrogen bonds, shape complementarity, and total score.

Across all designs that passed screening for ligand-water contacts, Rosetta design suggested the mutations K64(59)VAN, S96(92)V, E147(141)Y, and F165(159)AMVI (corresponding PYR1 numbering in parenthesis). Based on previous experience with poor prediction of electrostatic interactions by Rosetta, we excluded mutations at PYR1 positions 59 and 141, instead incorporating the PYR1^WIN^-mutation K59Q into designs. DNA sequences of the wild-type PYR1 sequence with combinations of the point mutations K59Q, S92V, F159A, F159V, A160I, and A160V were ordered (Integrated DNA Technologies) and cloned by Golden Gate Assembly ^51^ into the pND003 vector. Constructs were expressed in yeast surface display and binding affinity to WIN55212,2 was analyzed as previously described for isogenic variants.

## Supporting information

Complete data and primer list

## ASSOCIATED CONTENT

Supporting Information

The Supporting Information is available free of charge on the ACS Publications website.

Leonard_SI_2024.pdf

Supplemental_data&primers.xlsx

## AUTHOR INFORMATION

### Author Contributions

The manuscript was written by A.C.L., A.J.F., R.C., M.S., and T.A.W. with contributions from all other co-authors. The manuscript was approved by all co-authors.

## ACKNOWLEDGMENT

We would like to thank Sean Cutler and Ian Wheeldon at University of California, Riverside for helpful comments, and Kevin Kunstman and Stefan Green at Rush University for technical assistance with deep sequencing preparation. Simulations were run on Bridges2 and CU Boulder Alpine. A.C.L also thanks the NIH/CU Biophysics Training Program and the Interdisciplinary Quantitative Biology program at CU Boulder for ongoing support.

## ABBREVIATIONS

MD: molecular dynamics
H-bond: hydrogen bond

## REFERENCES

(1) Leonard, A. C.; Whitehead, T. A. Design and Engineering of Genetically Encoded Protein Biosensors for Small Molecules. Curr. Opin. Biotechnol. 2022, 78, 102787. 10.1016/j.copbio.2022.102787.

(2) Park, S. Y.; Peterson, F. C.; Mosquna, A.; Yao, J.; Volkman, B. F.; Cutler, S. R. Agrochemical Control of Plant Water Use Using Engineered Abscisic Acid Receptors. Nature 2015, 520 (7548), 545–548. 10.1038/nature14123.

(3) Wan, J.; Peng, W.; Li, X.; Qian, T.; Song, K.; Zeng, J.; Deng, F.; Hao, S.; Feng, J.; Zhang, P.; Zhang, Y.; Zou, J.; Pan, S.; Shin, M.; Venton, B. J.; Zhu, J. J.; Jing, M.; Xu, M.; Li, Y. A Genetically Encoded Sensor for Measuring Serotonin Dynamics. Nat. Neurosci. 2021, 24 (5), 746–752. 10.1038/s41593-021-00823-7.

(4) Wu, C. Y.; Roybal, K. T.; Puchner, E. M.; Onuffer, J.; Lim, W. A. Remote Control of Therapeutic T Cells through a Small Molecule-Gated Chimeric Receptor. Science 2015, 350 (6258), aab4077. 10.1126/SCIENCE.AAB4077.

(5) Duong, M. L. T.; Collinson-Pautz, M. R.; Morschl, E.; Lu, A.; Szymanski, S. P.; Zhang, M.; Brandt, M. E.; Chang, W. C.; Sharp, K. L.; Toler, S. M.; Slawin, K. M.; Foster, A. E.; Spencer, D. M.; Bayle, J. H. Two-Dimensional Regulation of CAR-T Cell Therapy with Orthogonal Switches. Molecular therapy oncolytics 2018, 12, 124–137. 10.1016/J.OMTO.2018.12.009.

(6) Giordano-Attianese, G.; Gainza, P.; Gray-Gaillard, E.; Cribioli, E.; Shui, S.; Kim, S.; Kwak, M. J.; Vollers, S.; Corria Osorio, A. D. J.; Reichenbach, P.; Bonet, J.; Oh, B. H.; Irving, M.; Coukos, G.; Correia, B. E. A Computationally Designed Chimeric Antigen Receptor Provides a Small-Molecule Safety Switch for T-Cell Therapy. Nature Biotechnology 2020 38:4 2020, 38 (4), 426–432. 10.1038/s41587-019-0403-9.

(7) Kvorjak, M.; Ruffo, E.; Tivon, Y.; So, V.; Parikh, A.; Deiters, A.; Lohmueller, J. Conditional Control of Universal CAR T Cells by Cleavable OFF-Switch Adaptors. ACS Synth. Biol. 2023, 12 (10), 2996–3007. 10.1021/acssynbio.3c00320.

(8) D’Ambrosio, V.; Pramanik, S.; Goroncy, K.; Jakočiūnas, T.; Schönauer, D.; Davari, M. D.; Schwaneberg, U.; Keasling, J. D.; Jensen, M. K. Directed Evolution of VanR Biosensor Specificity in Yeast. Biotechnology Notes 2020, 1, 9–15. 10.1016/j.biotno.2020.01.002.

(9) Taylor, N. D.; Garruss, A. S.; Moretti, R.; Chan, S.; Arbing, M. A.; Cascio, D.; Rogers, J. K.; Isaacs, F. J.; Kosuri, S.; Baker, D.; Fields, S.; Church, G. M.; Raman, S. Engineering an Allosteric Transcription Factor to Respond to New Ligands. Nat. Methods 2016, 13 (2), 177. 10.1038/NMETH.3696.

(10) Magnus, C. J.; Lee, P. H.; Bonaventura, J.; Zemla, R.; Gomez, J. L.; Ramirez, M. H.; Hu, X.; Galvan, A.; Basu, J.; Michaelides, M.; Sternson, S. M. Ultrapotent Chemogenetics for Research and Potential Clinical Applications. Science 2019, 364 (6436). 10.1126/SCIENCE.AAV5282/SUPPL_FILE/AAV5282S2.MOV.

(11) Weston, M.; Kaserer, T.; Wu, A.; Mouravlev, A.; Carpenter, J. C.; Snowball, A.; Knauss, S.; von Schimmelmann, M.; During, M. J.; Lignani, G.; Schorge, S.; Young, D.; Kullmann, D. M.; Lieb, A. Olanzapine: A Potent Agonist at the hM4D(Gi) DREADD Amenable to Clinical Translation of Chemogenetics. Sci Adv 2019, 5 (4), eaaw1567. 10.1126/sciadv.aaw1567.

(12) Herud-Sikimić, O.; Stiel, A. C.; Kolb, M.; Shanmugaratnam, S.; Berendzen, K. W.; Feldhaus, C.; Höcker, B.; Jürgens, G. A Biosensor for the Direct Visualization of Auxin. Nature 2021, 592 (7856), 768. 10.1038/S41586-021-03425-2.

(13) Beltrán, J.; Steiner, P. J.; Bedewitz, M.; Wei, S.; Peterson, F. C.; Li, Z.; Hughes, B. E.; Hartley, Z.; Robertson, N. R.; Medina-Cucurella, A. V.; Baumer, Z. T.; Leonard, A. C.; Park, S.-Y.; Volkman, B. F.; Nusinow, D. A.; Zhong, W.; Wheeldon, I.; Cutler, S. R.; Whitehead, T. A. Rapid Biosensor Development Using Plant Hormone Receptors as Reprogrammable Scaffolds. Nat. Biotechnol. 2022. 10.1038/s41587-022-01364-5.

(14) Cao, S.; Kang, S.; Mao, H.; Yao, J.; Gu, L.; Zheng, N. Defining Molecular Glues with a Dual-Nanobody Cannabidiol Sensor. Nat. Commun. 2022, 13 (1), 815. 10.1038/s41467-022-28507-1.

(15) Lee, G. R.; Pellock, S. J.; Norn, C.; Tischer, D.; Dauparas, J.; Anischenko, I.; Mercer, J. A. M.; Kang, A.; Bera, A.; Nguyen, H.; Goreshnik, I.; Vafeados, D.; Roullier, N.; Han, H. L.; Coventry, B.; Haddox, H. K.; Liu, D. R.; Yeh, A. H.-W.; Baker, D. Small-Molecule Binding and Sensing with a Designed Protein Family. bioRxiv 2023. 10.1101/2023.11.01.565201.

(16) Polizzi, N. F.; DeGrado, W. F. A Defined Structural Unit Enables de Novo Design of Small-Molecule-Binding Proteins. Science 2020, 369 (6508), 1227–1233. 10.1126/SCIENCE.ABB8330.

(17) Glasgow, A. A.; Huang, Y. M.; Mandell, D. J.; Thompson, M.; Ritterson, R.; Loshbaugh, A. L.; Pellegrino, J.; Krivacic, C.; Pache, R. A.; Barlow, K. A.; Ollikainen, N.; Jeon, D.; Kelly, M. J. S.; Fraser, J. S.; Kortemme, T. Computational Design of a Modular Protein Sense/response System. Science 2019, 366 (6468), 1024. 10.1126/SCIENCE.AAX8780.

(18) Polizzi, N. F.; Wu, Y.; Lemmin, T.; Maxwell, A. M.; Zhang, S.-Q.; Rawson, J.; Beratan, D. N.; Therien, M. J.; DeGrado, W. F. De Novo Design of a Hyperstable Non-Natural Protein-Ligand Complex with Sub-Å Accuracy. Nat. Chem. 2017, 9 (12), 1157–1164. 10.1038/nchem.2846.

(19) Shui, S.; Gainza, P.; Scheller, L.; Yang, C.; Kurumida, Y.; Rosset, S.; Georgeon, S.; Di Roberto, R. B.; Castellanos-Rueda, R.; Reddy, S. T.; Correia, B. E. A Rational Blueprint for the Design of Chemically-Controlled Protein Switches. Nature Communications 2021 12:1 2021, 12 (1), 1–12. 10.1038/s41467-021-25735-9.

(20) Kang, S.; Davidsen, K.; Gomez-Castillo, L.; Jiang, H.; Fu, X.; Li, Z.; Liang, Y.; Jahn, M.; Moussa, M.; DiMaio, F.; Gu, L. COMBINES-CID: An Efficient Method for De Novo Engineering of Highly Specific Chemically Induced Protein Dimerization Systems. J. Am. Chem. Soc. 2019, 141 (28), 10948–10952. 10.1021/jacs.9b03522.

(21) Tinberg, C. E.; Khare, S. D.; Dou, J.; Doyle, L.; Nelson, J. W.; Schena, A.; Jankowski, W.; Kalodimos, C. G.; Johnsson, K.; Stoddard, B. L.; Baker, D. Computational Design of Ligand Binding Proteins with High Affinity and Selectivity. Nature 2013, 501 (7466), 212. 10.1038/NATURE12443.

(22) Bade, R.; Chan, H.-F.; Reynisson, J. Characteristics of Known Drug Space. Natural Products, Their Derivatives and Synthetic Drugs. Eur. J. Med. Chem. 2010, 45 (12), 5646–5652. 10.1016/j.ejmech.2010.09.018.

(23) Boder, E. T.; Midelfort, K. S.; Wittrup, K. D. Directed Evolution of Antibody Fragments with Monovalent Femtomolar Antigen-Binding Affinity. Proc. Natl. Acad. Sci. U. S. A. 2000, 97 (20), 10701–10705. 10.1073/pnas.170297297.

(24) Hilvert, D. Critical Analysis of Antibody Catalysis. Annu. Rev. Biochem. 2000, 69, 751–793. 10.1146/annurev.biochem.69.1.751.

(25) Skerra, A. Alternative Binding Proteins: Anticalins - Harnessing the Structural Plasticity of the Lipocalin Ligand Pocket to Engineer Novel Binding Activities. FEBS J. 2008, 275 (11), 2677–2683. 10.1111/j.1742-4658.2008.06439.x.

(26) Flower, D. R.; North, A. C.; Sansom, C. E. The Lipocalin Protein Family: Structural and Sequence Overview. Biochim. Biophys. Acta 2000, 1482 (1-2), 9–24. 10.1016/s0167-4838(00)00148-5.

(27) Iyer, L. M.; Koonin, E. V.; Aravind, L. Adaptations of the Helix-Grip Fold for Ligand Binding and Catalysis in the START Domain Superfamily. Proteins 2001, 43 (2), 134–144. 10.1002/1097-0134(20010501)43:2<134::aid-prot1025>3.0.co;2-i.

(28) Létourneau, D.; Lefebvre, A.; Lavigne, P.; LeHoux, J.-G. The Binding Site Specificity of STARD4 Subfamily: Breaking the Cholesterol Paradigm. Mol. Cell. Endocrinol. 2015, 408, 53–61. 10.1016/j.mce.2014.12.016.

(29) Schrick, K.; Bruno, M.; Khosla, A.; Cox, P. N.; Marlatt, S. A.; Roque, R. A.; Nguyen, H. C.; He, C.; Snyder, M. P.; Singh, D.; Yadav, G. Shared Functions of Plant and Mammalian StAR-Related Lipid Transfer (START) Domains in Modulating Transcription Factor Activity. BMC Biol. 2014, 12, 70. 10.1186/s12915-014-0070-8.

(30) Mahtha, S. K.; Kumari, K.; Gaur, V.; Yadav, G. Cavity Architecture Based Modulation of Ligand Binding Tunnels in Plant START Domains. Comput. Struct. Biotechnol. J. 2023, 21, 3946–3963. 10.1016/j.csbj.2023.07.039.

(31) Park, S.-Y.; Qiu, J.; Wei, S.; Peterson, F. C.; Beltrán, J.; Medina-Cucurella, A. V.; Vaidya, A. S.; Xing, Z.; Volkman, B. F.; Nusinow, D. A.; Whitehead, T. A.; Wheeldon, I.; Cutler, S. R. An Orthogonalized PYR1-Based CID Module with Reprogrammable Ligand-Binding Specificity. Nat. Chem. Biol. 2024, 20 (1), 103–110. 10.1038/s41589-023-01447-7.

(32) Park, S.-Y.; Fung, P.; Nishimura, N.; Jensen, D. R.; Fujii, H.; Zhao, Y.; Lumba, S.; Santiago, J.; Rodrigues, A.; Chow, T.-F. F.; Alfred, S. E.; Bonetta, D.; Finkelstein, R.; Provart, N. J.; Desveaux, D.; Rodriguez, P. L.; McCourt, P.; Zhu, J.-K.; Schroeder, J. I.; Volkman, B. F.; Cutler, S. R. Abscisic Acid Inhibits Type 2C Protein Phosphatases via the PYR/PYL Family of START Proteins. Science 2009, 324 (5930), 1068–1071. 10.1126/science.1173041.

(33) Lumba, S.; Cutler, S.; McCourt, P. Plant Nuclear Hormone Receptors: A Role for Small Molecules in Protein-Protein Interactions. Annu. Rev. Cell Dev. Biol. 2010, 26, 445–469. 10.1146/annurev-cellbio-100109-103956.

(34) Steiner, P. J.; Swift, S. D.; Bedewitz, M.; Wheeldon, I.; Cutler, S. R.; Nusinow, D. A.; Whitehead, T. A. A Closed Form Model for Molecular Ratchet-Type Chemically Induced Dimerization Modules. Biochemistry 2023, 62 (2), 281–291. 10.1021/acs.biochem.2c00172.

(35) Melcher, K.; Ng, L. M.; Zhou, X. E.; Soon, F. F.; Xu, Y.; Suino-Powell, K. M.; Park, S. Y.; Weiner, J. J.; Fujii, H.; Chinnusamy, V.; Kovach, A.; Li, J.; Wang, Y.; Li, J.; Peterson, F. C.; Jensen, D. R.; Yong, E. L.; Volkman, B. F.; Cutler, S. R.; Zhu, J. K.; Xu, H. E. A Gate–latch–lock Mechanism for Hormone Signalling by Abscisic Acid Receptors. Nature 2009, 462 (7273), 602–608. 10.1038/nature08613.

(36) Zhao, C.; Shukla, D. Molecular Basis of the Activation and Dissociation of Dimeric PYL2 Receptor in Abscisic Acid Signaling. Phys. Chem. Chem. Phys. 2022, 24 (2), 724–734. 10.1039/d1cp03307g.

(37) Kim, S.; Chen, J.; Cheng, T.; Gindulyte, A.; He, J.; He, S.; Li, Q.; Shoemaker, B. A.; Thiessen, P. A.; Yu, B.; Zaslavsky, L.; Zhang, J.; Bolton, E. E. PubChem 2023 Update. Nucleic Acids Res. 2023, 51 (D1), D1373–D1380. 10.1093/nar/gkac956.

(38) Petersen, B. M.; Kirby, M.; Chrispens, K.; Irvin, O.; Strawn, I.; Haas, C.; Walker, A.; Baumer, Z. T.; Ulmer, S.; Ayala, E.; Rhodes, E. R.; Guthmiller, J.; Steiner, P. J.; Whitehead, T. A. An Integrated Technology for Quantitative Wide Mutational Scanning of Human Antibody Fab Libraries. bioRxiv, 2024, 2024.01.16.575852. 10.1101/2024.01.16.575852.

(39) Tirado-Rives, J.; Jorgensen, W. L. Contribution of Conformer Focusing to the Uncertainty in Predicting Free Energies for Protein-Ligand Binding. J. Med. Chem. 2006, 49 (20), 5880–5884. 10.1021/jm060763i.

(40) Whitehead, T. A.; Chevalier, A.; Song, Y.; Dreyfus, C.; Fleishman, S. J.; De Mattos, C.; Myers, C. A.; Kamisetty, H.; Blair, P.; Wilson, I. A.; Baker, D. Optimization of Affinity, Specificity and Function of Designed Influenza Inhibitors Using Deep Sequencing. Nat. Biotechnol. 2012, 30 (6), 543–548. 10.1038/nbt.2214.

(41) Park, Y.; Metzger, B. P. H.; Thornton, J. W. The Simplicity of Protein Sequence-Function Relationships. bioRxiv 2024. 10.1101/2023.09.02.556057.

(42) Maguire, J. B.; Haddox, H. K.; Strickland, D.; Halabiya, S. F.; Coventry, B.; Griffin, J. R.; Pulavarti, S. V. S. R. K.; Cummins, M.; Thieker, D. F.; Klavins, E.; Szyperski, T.; DiMaio, F.; Baker, D.; Kuhlman, B. Perturbing the Energy Landscape for Improved Packing during Computational Protein Design. Proteins 2021, 89 (4), 436–449. 10.1002/prot.26030.

(43) Wei, B.; Zhang, Y.; Gong, X. DeepLPI: A Novel Deep Learning-Based Model for Protein-Ligand Interaction Prediction for Drug Repurposing. Sci. Rep. 2022, 12 (1), 18200. 10.1038/s41598-022-23014-1.

(44) Jin, Z.; Wu, T.; Chen, T.; Pan, D.; Wang, X.; Xie, J.; Quan, L.; Lyu, Q. CAPLA: Improved Prediction of Protein-Ligand Binding Affinity by a Deep Learning Approach Based on a Cross-Attention Mechanism. Bioinformatics 2023, 39 (2). 10.1093/bioinformatics/btad049.

(45) Wang, D. D.; Wu, W.; Wang, R. Structure-Based, Deep-Learning Models for Protein-Ligand Binding Affinity Prediction. J. Cheminform. 2024, 16 (1), 2. 10.1186/s13321-023-00795-9.

(46) Rezaei, M. A.; Li, Y.; Wu, D.; Li, X.; Li, C. Deep Learning in Drug Design: Protein-Ligand Binding Affinity Prediction. IEEE/ACM Trans. Comput. Biol. Bioinform. 2022, 19 (1), 407–417. 10.1109/TCBB.2020.3046945.

(47) Dauparas, J.; Lee, G. R.; Pecoraro, R.; An, L.; Anishchenko, I.; Glasscock, C.; Baker, D. Atomic Context-Conditioned Protein Sequence Design Using LigandMPNN. bioRxiv, 2023, 2023.12.22.573103. 10.1101/2023.12.22.573103.

(48) Zhang, Z.; Shen, W.; Liu, Q.; Zitnik, M. PocketGen: Generating Full-Atom Ligand-Binding Protein Pockets. bioRxiv, 2024, 2024.02.25.581968. 10.1101/2024.02.25.581968.

(49) Wrenbeck, E. E.; Klesmith, J. R.; Stapleton, J. A.; Adeniran, A.; Tyo, K. E. J.; Whitehead, T. A. Plasmid-Based One-Pot Saturation Mutagenesis. Nat. Methods 2016, 13 (11), 928–930. 10.1038/nmeth.4029.

(50) Steiner, P. J.; Bedewitz, M. A.; Medina-Cucurella, A. V.; Cutler, S. R.; Whitehead, T. A. A Yeast Surface Display Platform for Plant Hormone Receptors: Toward Directed Evolution of New Biosensors. AIChE J. 2020, 66 (3). 10.1002/aic.16767.

(51) Daffern, N.; Francino-Urdaniz, I.; Baumer, Z. T.; Whitehead, T. A. Benchmarking Cassette-Based Deep Mutagenesis by Golden Gate Assembly. bioRxiv, 2023, 2023.04.13.536781. 10.1101/2023.04.13.536781.

