## Supplemental figures for "Rationalizing diverse binding mechanisms to the same protein fold: insights for ligand recognition and biosensor design"

### Supplementary Figures

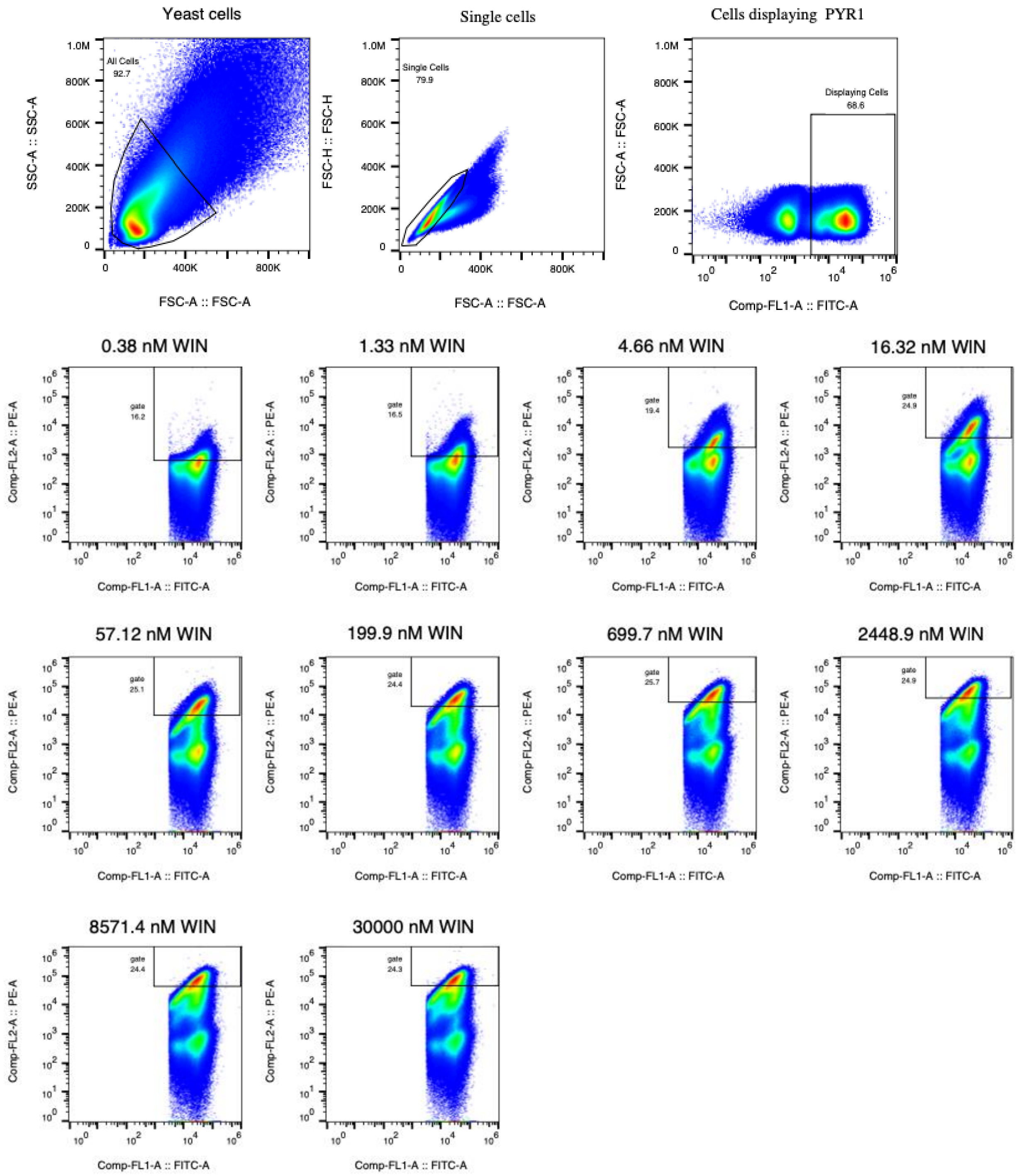


**Figure S1.** Cytograms demonstrating cell sorting procedure for deep mutational scanning analysis, showing PYR1$^{WIN}$ replicate A as replicate cytograms. Gates have been redrawn in FlowJo software, consult the supplemental data spreadsheet for precise gating statistics for each sensor replicate.

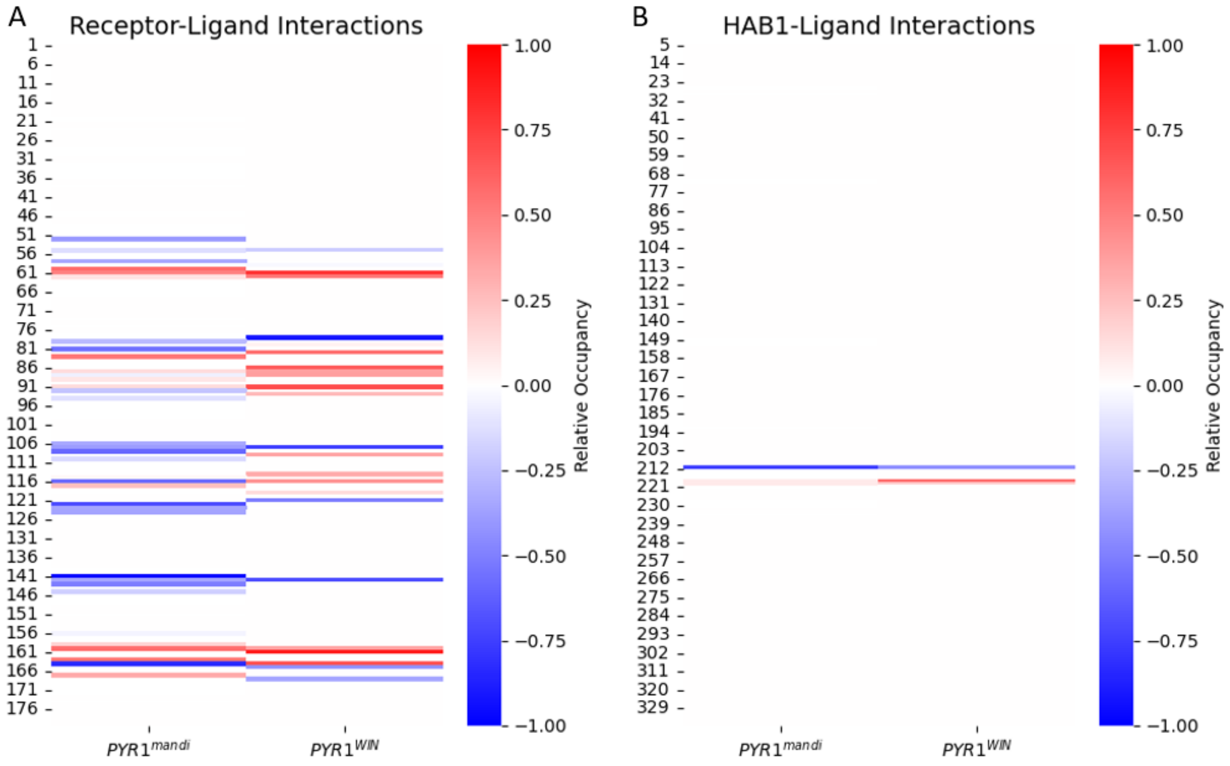


**Figure S2:** **Numerous water-mediated interactions support ligand binding.** These heatmaps display the relative occupancy of water-mediated vs direct non-bonded interactions between each protein residue and the ligand. A value of $R \simeq 0$ would mean that either a non-bonded interaction is not frequently formed between this residue and the ligand or that the interaction is formed equally as a direct and water-mediated interaction. A value of $R \simeq -1$ would mean that there is a water-mediated interaction formed for close to 100% of all trajectories and that there is no significant period of the trajectory for which a bonded interaction is formed between this same residue and the ligand. (A) There are numerous residues in the receptor for which $R<-0.7$ in both Mandipropamid and WIN. For $PYR1^{mandi}$ these residues are … For $PYR1^{WIN}$ these residues are … For $PYR1^{WIN}$, residue W385 has an R value of -0.45 which is lower than the threshold established, but this residue was still included for further analysis of the water-mediated h-bond was present for >95% of trajectories and this was previously established to be an important water-mediated interaction. It is important to note that no other residues had an R score between 0.45 and 0.7 and also formed water mediated interactions for >95% of trajectories.

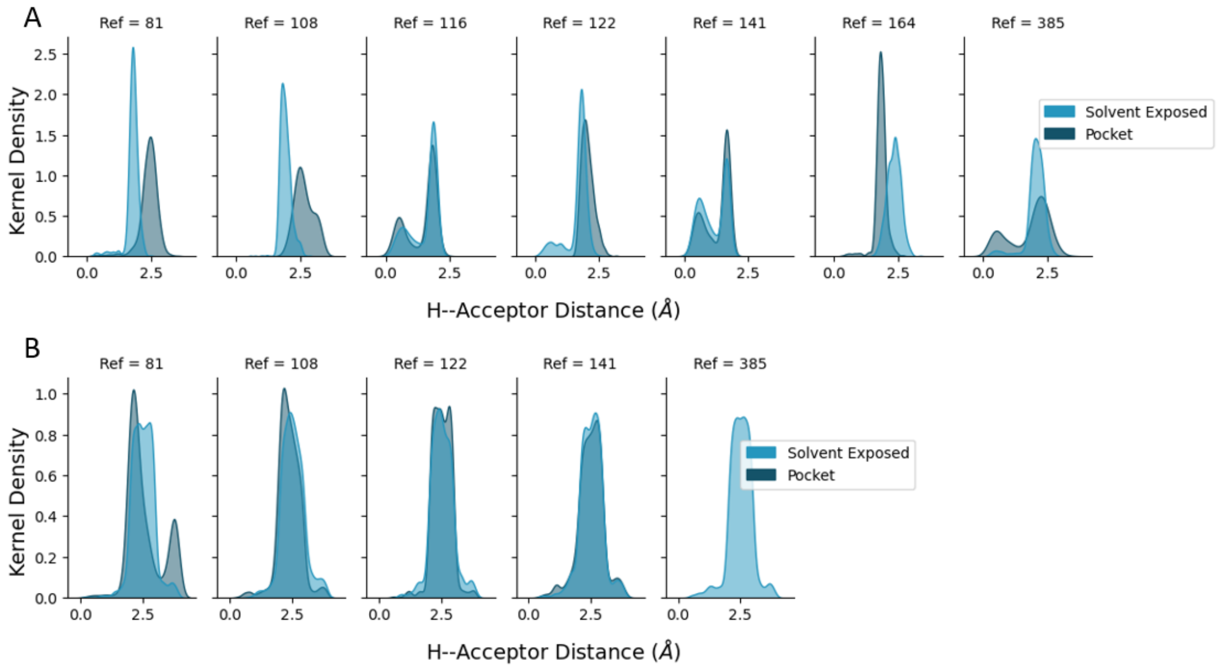


**Figure S3:** **A standard strong and stable h-bond establishes thresholds for water-mediated h-bonds.** We have included control residues with the same residue type as each residue that was determined to have a primarily water-mediated h-bond with the ligand (reference residue). The solvent exposed control is a residue as the same type as the reference residue but located in a solvent exposed position on the receptor. The pocket control resid is a residue the same type as the reference and located within the binding pocket, but with a combined <10% contact with the ligand from either direct or water-mediated interactions. (A) The control residues from PYR1$^{mandi}$ have a combined mean H–acceptor distance of 1.81 Å and 93% of the control bonds have a mean distance < 2.5 Å. Also, 80% of these residues have a standard deviation < 0.45 Å. (B) The control residues for PYR1$^{WIN}$ have a combined mean H–acceptor distance of 2.42 Å and 89% of the control bonds have a mean distance < 2.5 Å. Also, 89% of these residues have a standard deviation < 0.45 Å. This data allows increased confidence in the thresholds established for use dividing the water mediated h-bonds into those which are strong and stable and those which are not.

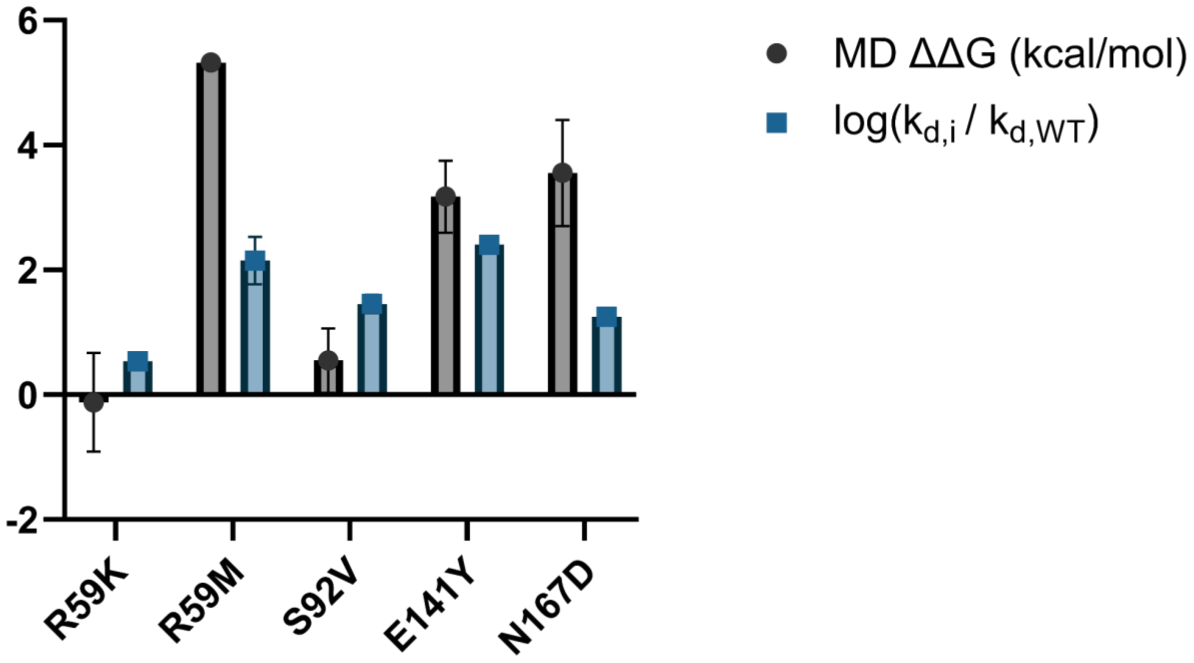


**Figure S4.** Comparison of ΔΔG calculated through MD simulations to experimentally-determined $log(\frac{K_{d,eff,i}}{K_{d,eff,WT}})$ for five chosen mutations to the PYR1$^{mandi}$ sensor. MD ΔΔG calculated from relative free energy calculations performed with Hamiltonian replica exchange simulations, in units of kcal/mol. The calculated $log(\frac{K_{d,eff,i}}{K_{d,eff,WT}})$ for each mutation is reported in main text figure 3, with exact values available in the supplemental data file.

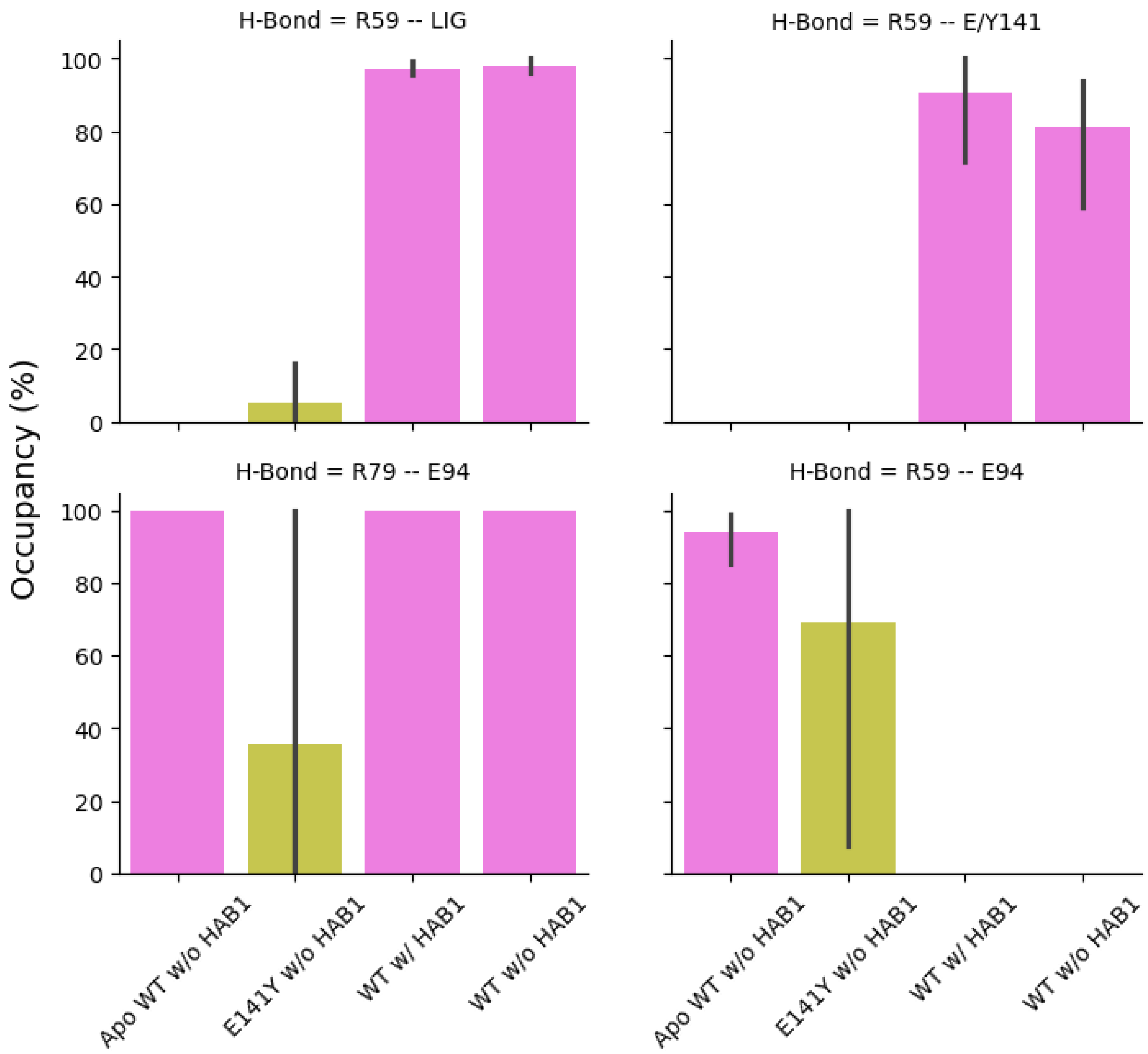


**Figure S5:** **Salt bridge formation is significantly impacted by mutation.** The formation of a salt bridge between R59 and the ligand appears to be central to the stabilization of the ligand within the binding pocket prior to HAB1 secondary binding. The R59–ligand salt bridge is disrupted by the E141Y mutation to PYR1$^{mandi}$. The absence of the R59–ligand salt bridge is strongly correlated to the absence of the R59-E/Y141 salt bridge which is disrupted directly by the E141Y mutation. In the absence of a salt-bridge between R59-E/Y141 R59 instead forms a salt-bridge with E94 which is also consistent with the salt-bridge structure found in Apo PYR1$^{mandi}$.

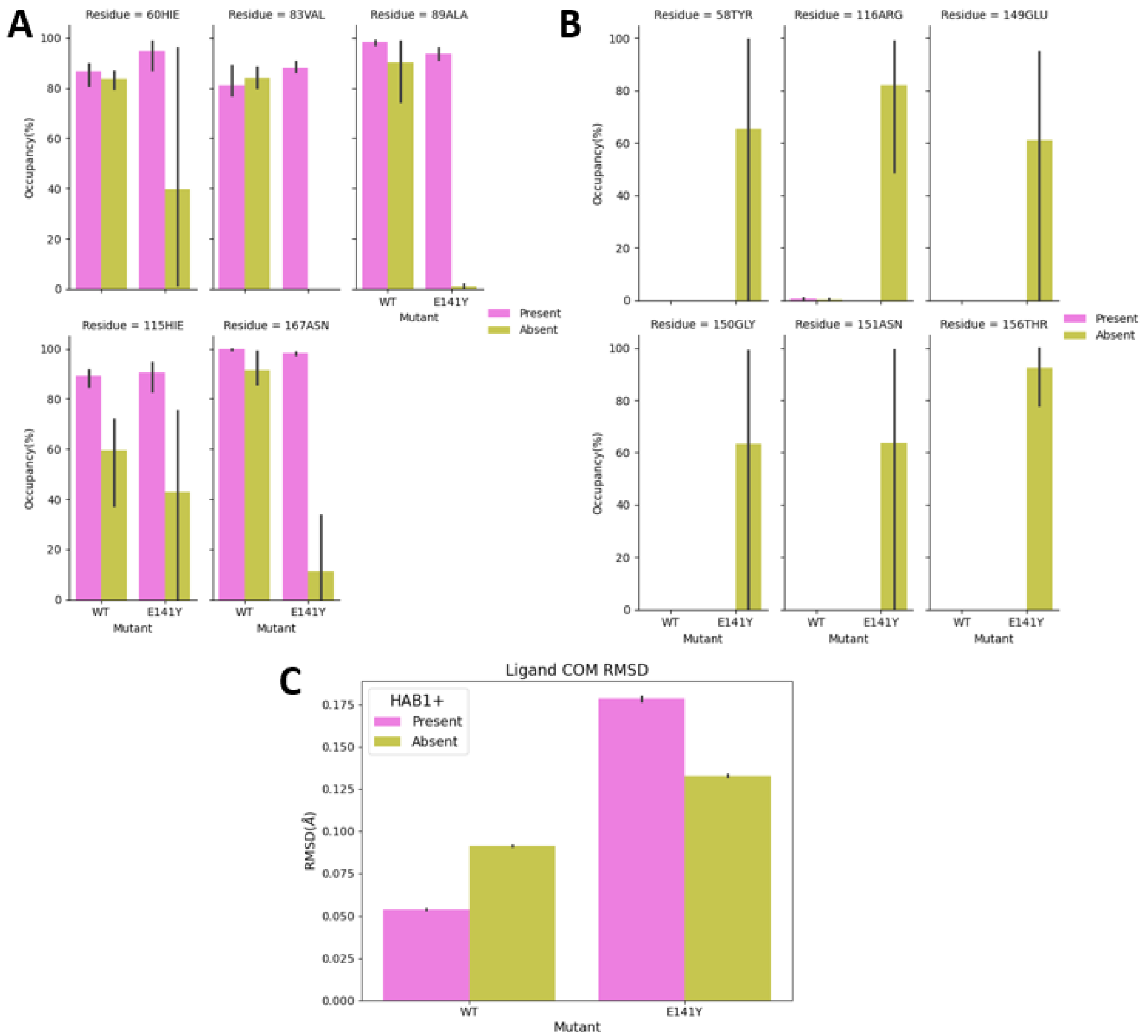


**Figure S6: E141Y mutation leads the ligand to adopt a significantly different conformation.** We examine additional ligand – protein non-bonded interactions to quantify the difference between the ligand binding location PYR1$^{mandi/E141Y}$ vs PYR1$^{mandi}$. (A) We plot the non-bonded interactions which form in >90% of trajectories for the PYR1$^{mandi}$ complex. We observe that all of these interactions are significantly disrupted for PYR1$^{mandi/E141Y}$ in the absence of HAB1. (B) Additionally we observe new distinct non-bonded interactions forming with the protein in it's new binding location. (C ) Regardless of whether PYR1$^{mandi/E141Y}$ is in complex with HAB1, the ligand is significantly destabilized as observed by the significant increase in center of mass (COM) RMSD.

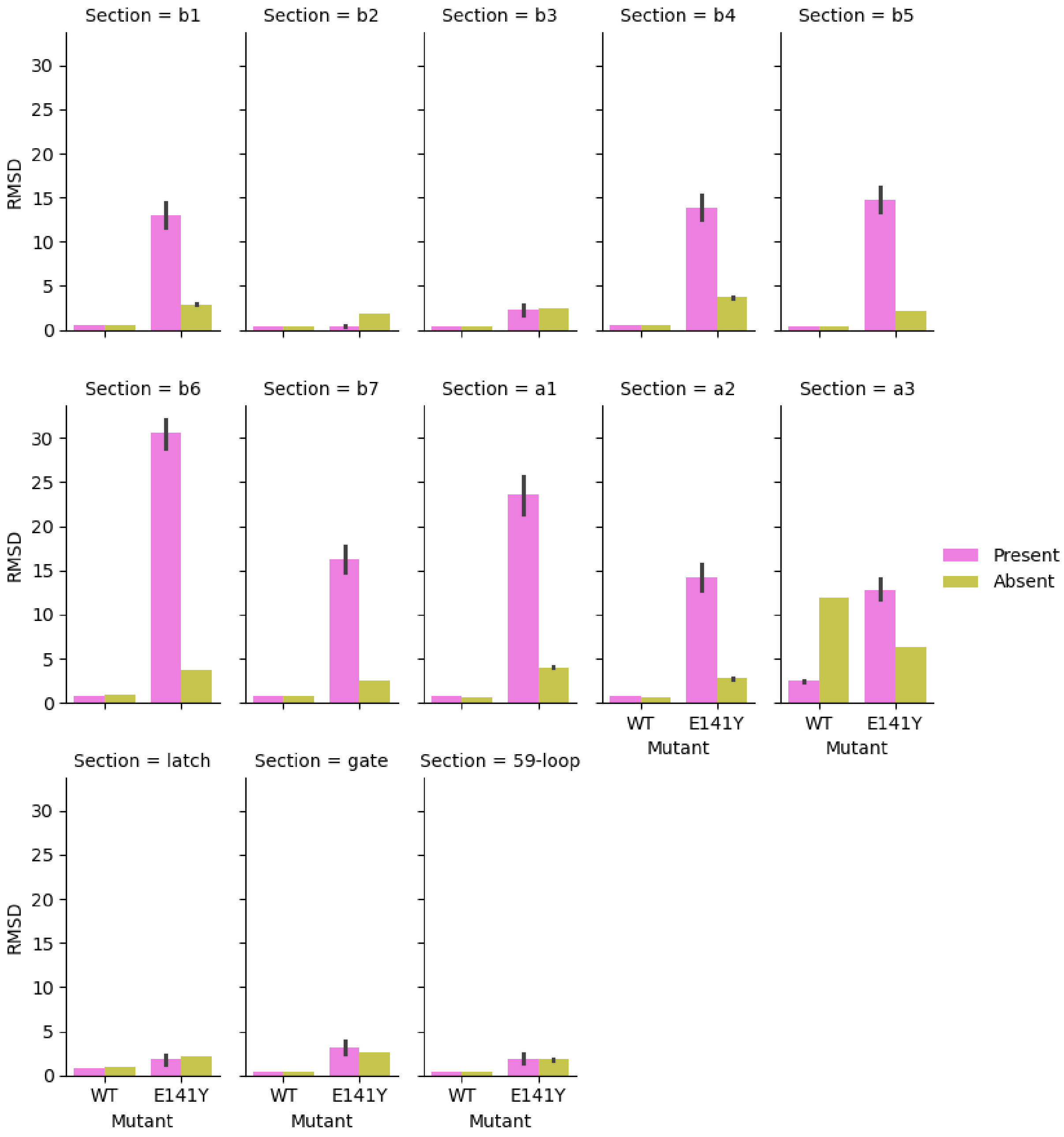


**Figure S7:** **E141Y mutation leads to minor distortion of the receptor.** Here we compare heavy atom RMSD for all variants relative to the centroid structure for WT PYR1 in complex with HAB1. Significantly increased RMSD values suggest that the structure is either significantly more flexible or exhibits a distinct protein conformation. The only variant which consistently exhibits a distinct conformation is PYR1$^{mandi/E141Y}$ in complex with HAB1 likely because the protein–protein binding interface prevented the ligand from reorienting to a more energetically favorable conformation thus forcing the ligand to distort the protein structure to reach a metastable conformation.

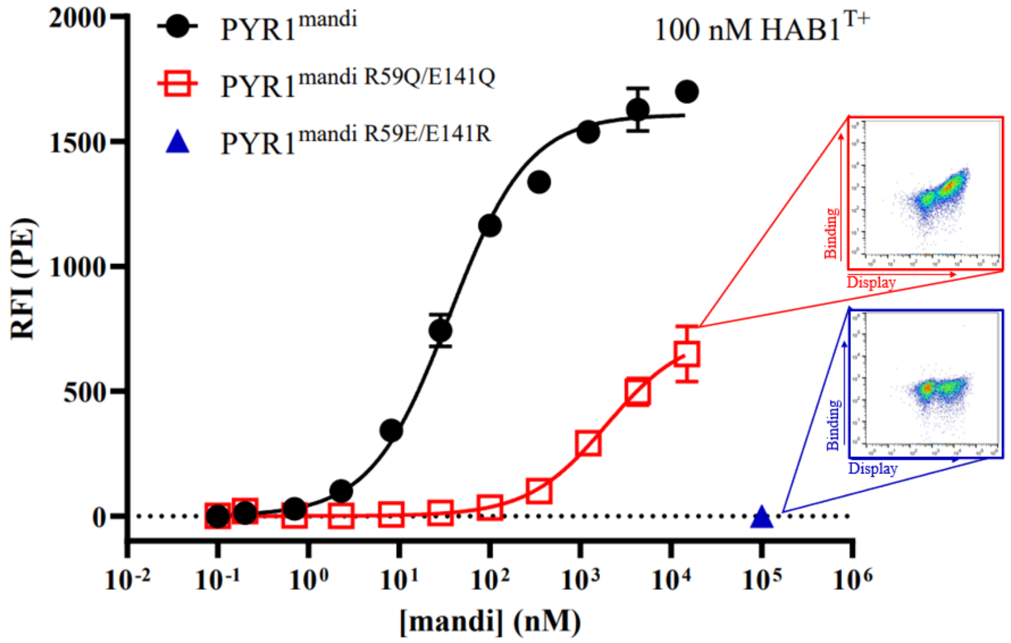


**Figure S8.** Biological replicate yeast surface display titration for PYR1$^{mandi}$, PYR1$^{mandi\ R59Q/E141Q}$, and PYR1$^{mandi\ R59E/E141R}$. Yeast surface display titrations of PYR1$^{mandi}$, PYR1$^{mandi\ R59Q/E141Q}$, and PYR1$^{mandi\ R59E/E141R}$. Relative fluorescence intensity (RFI) is determined by secondary labeling using streptavidin-phycoerythrin after initial labeling with indicated ligand concentration and 100 nM biotinylated HAB1$^{T+}$. Error bars represent 1 s.d. In relative fluorescence for n = 2 replicates. Inset cytograms show binding of HAB1$^{T+}$ versus display of PYR1 variant on the surface of yeast. The cytogram for PYR1$^{mandi\ R59Q/E141Q}$ shows the positive result of binding and the cytogram of PYR1$^{mandi\ R59E/E141R}$ shows the lack of binding for that variant.

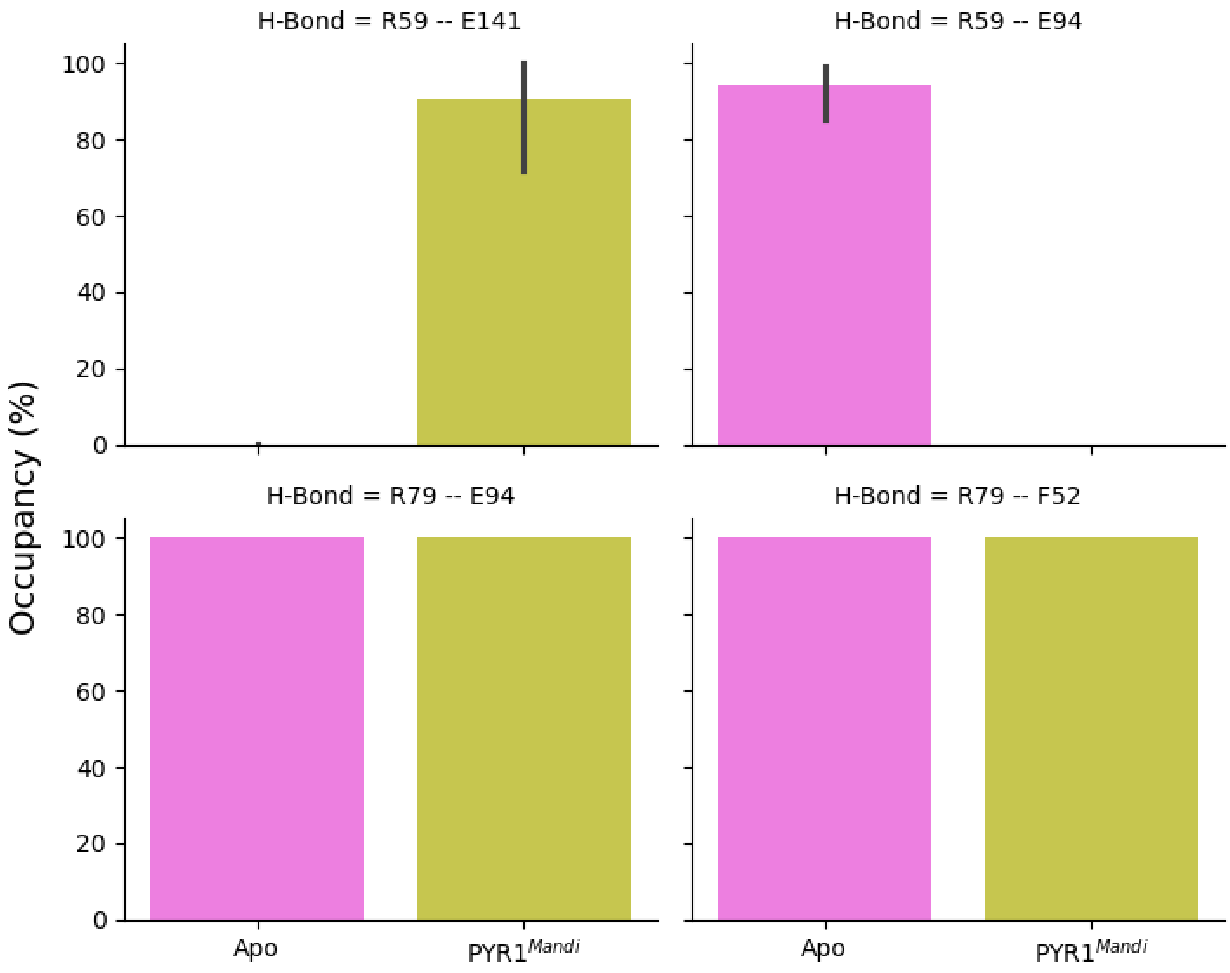


**Figure S9:** The Apo structure of the binding pocket is defined by stable salt bridges R59 – E94 and R79 – E94 as well as the h-bond R79 – F52 formed with the backbone of F52.

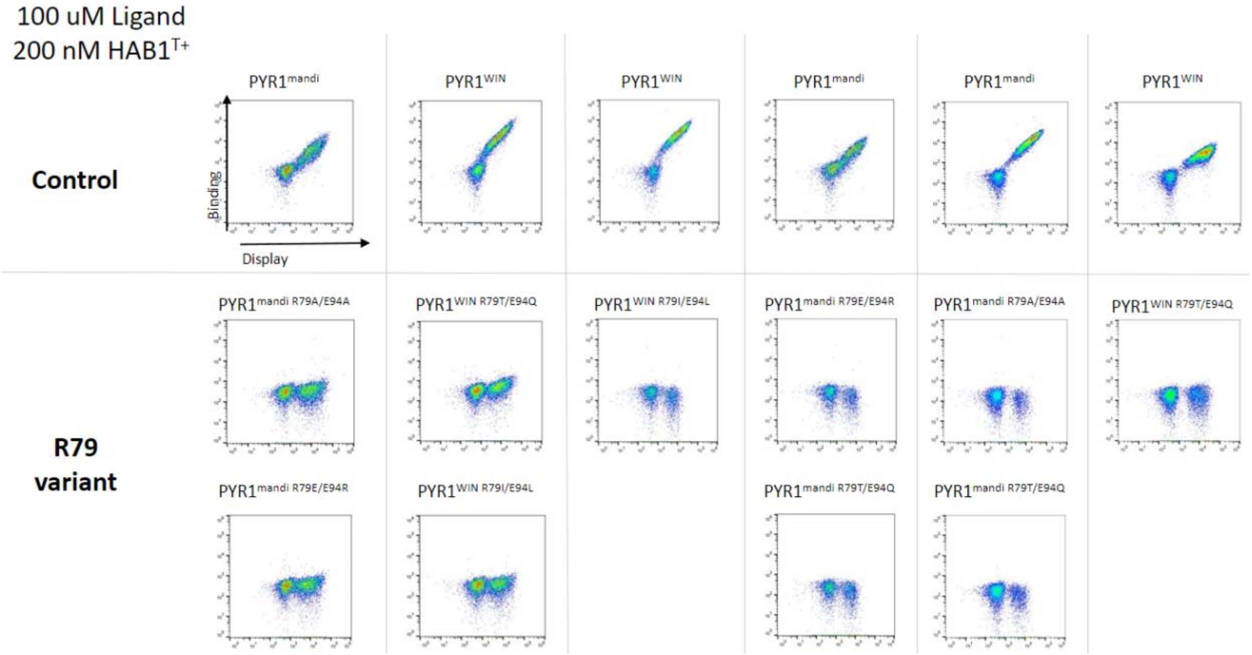


**Figure S10.** Cytograms of PYR1 R79-mutant binding to either 100μM mandipropamid or WIN55,212-2, as indicated, and 200nM HAB1$^{T+}$. Each cytogram shows a single experimental replicate, experiments run on different days are separated by vertical lines and grouped with the corresponding control.

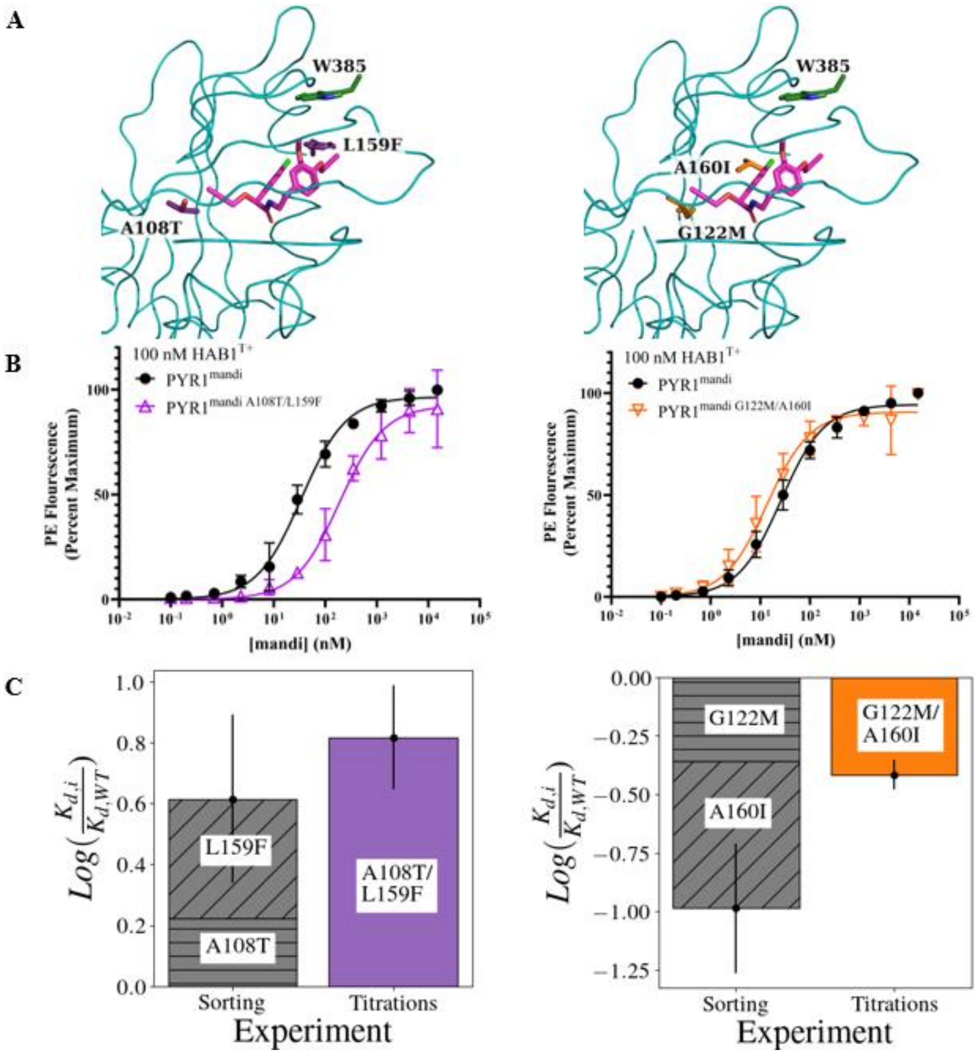


**Figure S11:** **Mutational additivity or non-additivity indicate conformer selection in engineered biosensors.** **A**) Structures of PYR1$^{\text{mandi A108T/L159F}}$ (left) and PYR1$^{\text{mandi A160I/G122M}}$ (right). The PYR1 backbone is shown as a teal ribbon. Ligands and mutated residues are shown as sticks. B) Yeast surface display titrations of multi-mutant biosensors compared with original sensor. PE fluorescence is determined by secondary labeling using streptavidin-phycoerythrin after initial labeling with indicated ligand concentration and 100 nM biotinylated HAB1$^{T+}$. Error bars represent 1 s.d. in relative PE fluorescence for n=4 (two technical replicates for two biological replicates). C) Predicted additive improvement of mutations in binding affinity compared to experimentally observed relative binding affinity. Error bars represent the summation of mean absolute error for each mutation (grey) or mean absolute error for titrations (colored).

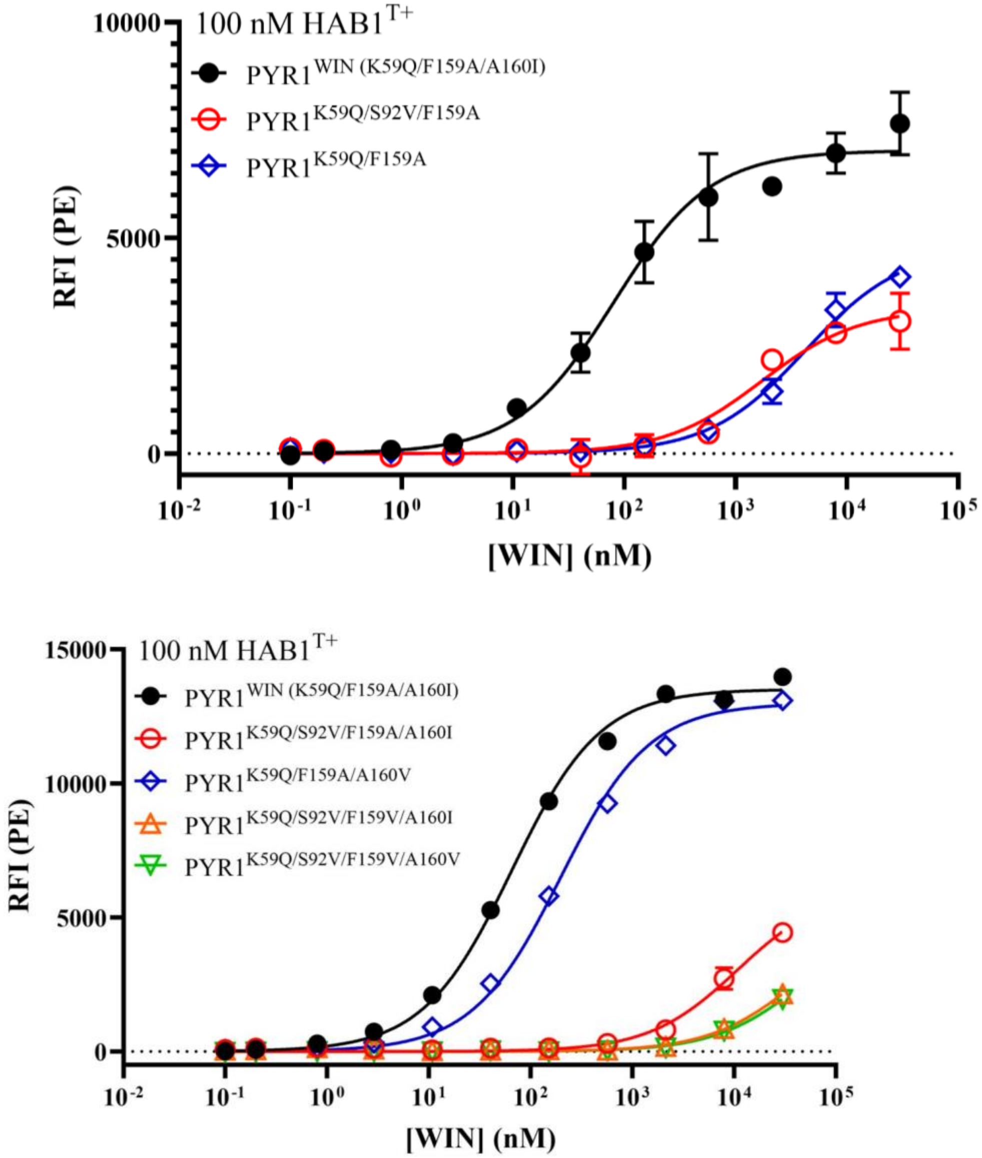


**Figure S12.** replicate of PYR1$^{WIN}$, PYR1$^{K59Q/S92V/F159A}$, and PYR1$^{K59Q//F159A}$ titration curves plus other variants Yeast surface display titrations of computationally designed PYR1 biosensors binding WIN55,212-2, compared wild-type control PYR1$^{WIN}$ sensor shown in black. PE fluorescence is determined by secondary labeling using streptavidin-phycoerythrin after initial labeling with indicated ligand concentration and 100 nM biotinylated HAB1$^{T+}$. Error bars represent 1 s.d. in relative PE fluorescence for n=2 (two technical replicates for one biological replicate).

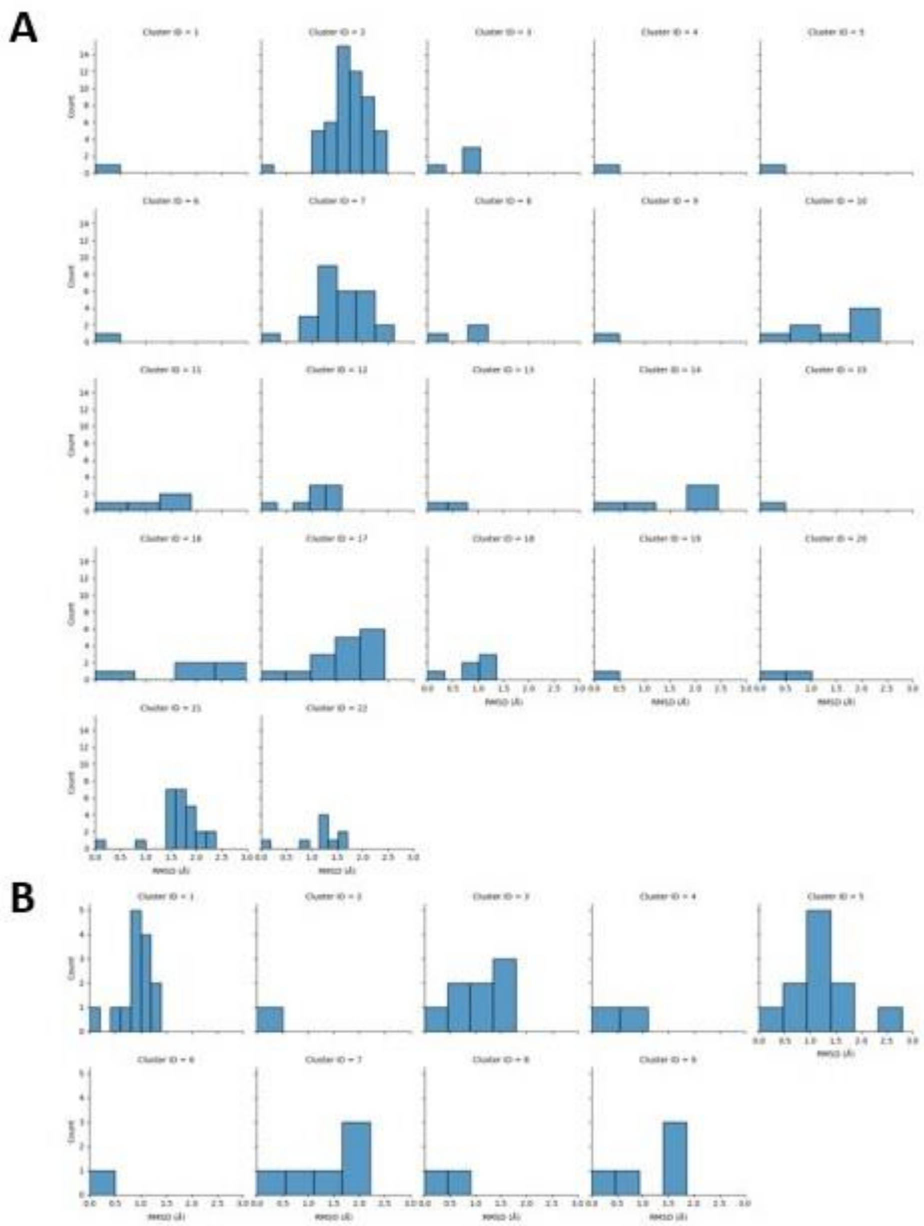

**Figure S13:** We examined the diversity in structures within the defined ligand clusters for Mandipropamid (A) and WIN (B) and we found that the clusters have much lower structural diversity than the non-clustered structures with a mean RMSD of 1-1.5Å compared to 3.75Å and 2.5Å without clustering for Mandipropamid and WIN respectively.
